# A bipartite mechanism for condensin II activation in mitosis

**DOI:** 10.64898/2026.02.02.703380

**Authors:** Damla Tetiker, Kumiko Samejima, Yan Li, David Schaumann, David Barford, Luis Aragon, William C. Earnshaw, Erin E. Cutts, Eugene Kim, Kyle W. Muir

**Affiliations:** Max Planck Institute of Biophysics, 60438, Frankfurt am Main, Germany; IMPRS on Cellular Biophysics, Max-von-Laue-Straße 3, 60438, Frankfurt am Main, Germany; Institute of Cell Biology, University of Edinburgh, Edinburgh, UK; MRC Laboratory of Molecular Biology, Francis Crick Avenue, Cambridge, CB2 0QH, UK; DNA Motors Group, MRC Laboratory of Medical Sciences (LMS), Du Cane Road, London, W12 0HS, UK; School of Biosciences, Faculty of Science, University of Sheffield, UK; Nucleic Acid Institute, University of Sheffield, UK

## Abstract

Human condensin II is a constitutively nuclear molecular motor that initiates chromosome organization in early mitosis. How condensin II is activated specifically in mitosis remains unknown. Here, we describe the molecular mechanism underlying condensin II auto-repression and activation. By determining multiple structural states of condensin II, we discovered that an autoinhibitory tail within the NCAPD3 subunit (NCAPD3^Tail^) holds the complex in a conformation that is incompatible with DNA capture. Deletion of NCAPD3^Tail^ spontaneously activates condensin II in cells, illuminating its autoinhibitory role *in vivo*. We further show this translates to increased loop DNA formation by condensin II *in vitro*. Direct competition for the NCAPD3^Tail^ binding site on condensin II enables the putative activator protein M18BP1 to liberate a key DNA-binding element in the NCAPH2 N-terminus, enabling DNA capture. Unexpectedly, M18BP1 not only relieves autoinhibition but also directly contributes to DNA organization by forming a positively charged loop that enhances DNA-anchoring by condensin II. Together, these findings reveal a bipartite activation mechanism wherein M18BP1 relieves autorepression, and renders condensin II biochemically competent to form stable DNA loops, ensuring highly stringent regulation of mitotic chromosome formation.

## Introduction

During mitosis, chromosomes undergo a dramatic reorganization from their ‘diffuse’, physically entangled interphasic architecture into discrete resolved sister chromatids ^1,2 3^. The disentanglement process is fundamental to ensure that sister chromatids can be mechanically separated and evenly divided between daughter cells following the dissolution of sister-chromatid cohesion at the metaphase-to-anaphase transition^4-6^.

Mitotic chromosome assembly is largely driven by condensins, that belong to the structural maintenance of chromosomes (SMC) family of DNA-binding ATPase motors ^7,8^. A prevailing hypothesis is that condensins reshape chromatin through ATP-dependent DNA loop extrusion in which DNA is actively reeled into growing loops, together with DNA-bridging activities that stabilize long-range contacts ^9-12^.

Condensins are pentameric complexes, in which a single ‘kleisin’ protein, links together the two accessory HEAT-repeat, HAWK, proteins around a heterodimeric SMC-ATPase core ^7^. Most eukaryotes, including humans, possess two condensin paralogues, condensin I, and condensin II ^8,13^, which share the same SMC2/4 subunits but differ in the non-SMC subunits: Kleisin (NCAPH and NCAPH2 for Condensin I and II respectively) and two HEAT-repeat HAWK subunits (NCAPG/D2 and NCAPG2/D3 for Condensin I and II respectively) ^8^. Condensin II is present in the nucleus throughout the cell cycle and initiates chromosome condensation at mitotic entry, whereas condensin I gains access to chromosomes only after nuclear envelope breakdown and forms loops nested within larger condensin II–dependent loops ^1,3,14,15^.

Despite its central role in shaping mitotic chromosomes, how condensin II activity is temporally restricted to mitosis remains poorly understood. Condensin II is an active ATPase ^16-18^ capable of binding and organizing DNA into loops ^19^, yet it does not globally compact chromatin during interphase. Condensin II is proposed to be inhibited during interphase through repression by a putative inhibitor MCPH1^17^. Genetic perturbations further suggest intrinsic autoinhibitory elements within the HAWK subunits ^20,21^. Activation at mitotic entry has been proposed to require condensin II phosphorylation^22-24^, and binding to a mitotic regulator, M18BP1^16,25^..Although a variety of regulatory factors, post-translational modifications, and potential autoinhibitory elements have been implicated in controlling condensin II, how condensin II is selectively activated at mitotic entry remains a fundamentally unsolved question.

Here, we employ a series of single-molecule, structural biochemistry and cell biological experiments to reveal the molecular basis of condensin II activation and activity during mitosis. This revealed an autoinhibitory mechanism in which the C-terminal NCAPD3^Tail^ sequesters the NCAPH2^Neck^ domain, preventing the complex from engaging chromatin. We found that phosphorylation of the regulator M18BP1 by CDK1 substantially enhances its ability to activate condensin II, which it achieves by releasing the NCAPD3^Tail^. Moreover, we observed that M18BP1 not only overcomes autoinhibition, but structurally augments a DNA loop anchor within condensin II to enable stable DNA loop formation. Overall this provides a molecular rationale for how condensin II activity is restricted during interphase, and activated in a CDK1-dependent manner during mitosis by a bipartite mechanism that relieves autoinhibition, whilst conferring substantially enhanced DNA organizing properties.

## Results

### Structure of apo condensin II

To investigate the molecular architecture of condensin II, we first determined the structure of the apo complex, in the absence of ATP and DNA, using cryo-electron microscopy (cryo-EM) to an overall resolution of 3.6 Å (Figure S1., Figure S2A-F). In contrast to structures of other eukaryotic SMCs such as cohesin and budding yeast condensin I, condensin II assembles a dimer, as well as a monomeric pentamer. For further analysis we performed particle-symmetry expansion and determined a monomer reconstruction at a resolution of 3.6 Å (Figure 1B., Figure S1B,C, Figure S2A-F).

**Figure 1:**
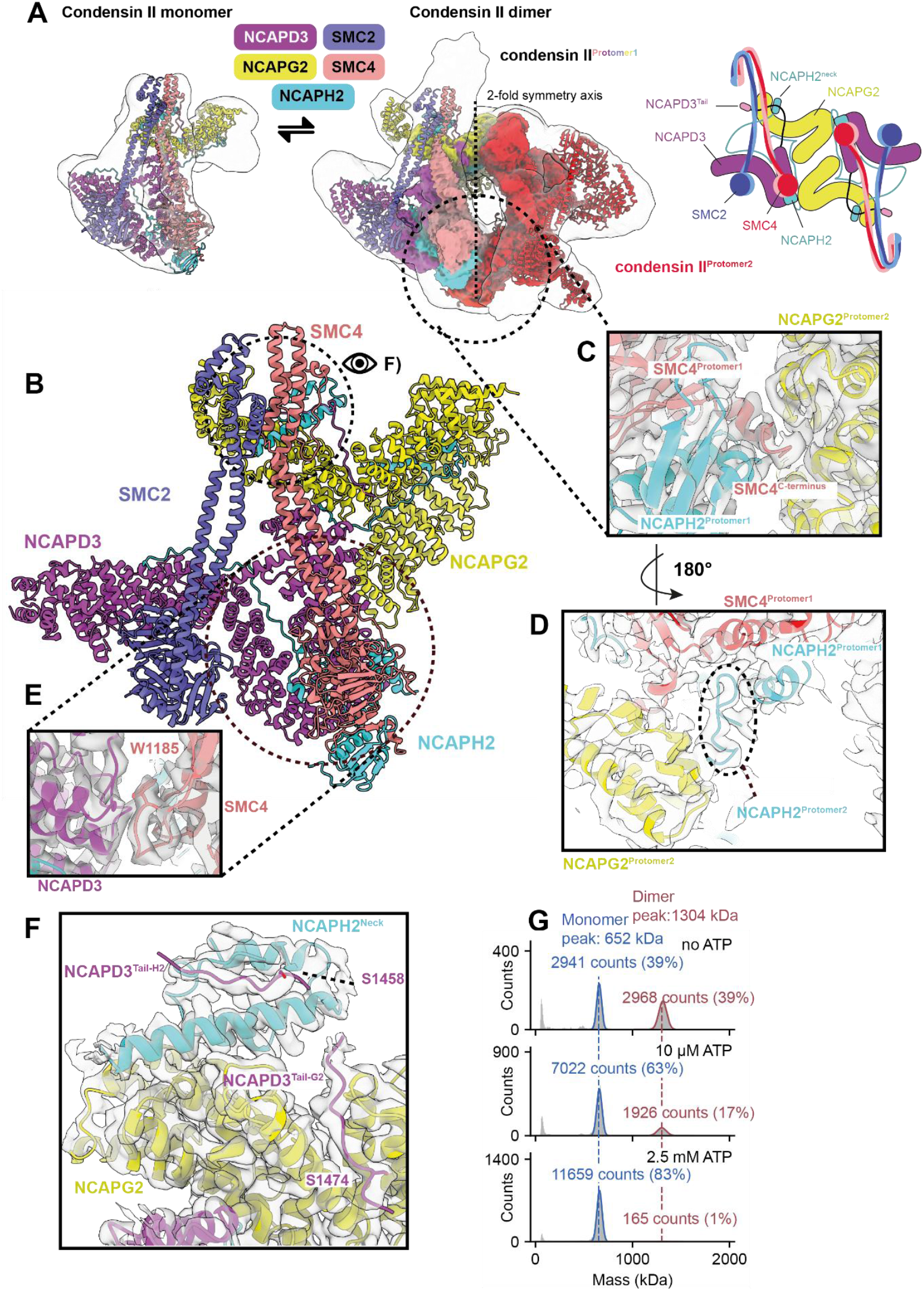
Apo condensin II is in a monomer-dimer equilibrium and forms DNA loops as a monomer. **(A)** Cryo-EM 3D reconstructions of apo condensin II in a monomeric and dimeric state. Cartoon schematic of dimeric condensin II **(B)** Overall structural model of an apo condensin II protomer monomer. **(C)** Details of the dimerization interface of condensin II through the C-terminus of SMC4^Protomer 1^ and NCAPG2^Protomer 2^. **(D)** Cryo-EM density displayed over dimerization interface mediated by contacts between SMC4^Protomer1^ and NCAPH2^Protomer1^, and NCAPH2^Protomer2^. **(E)** Cryo-EM density shows NCAPD3 contacts SMC4 through the W-loop of its ATPase domain. **(F)** The NCAPD3^Tail^ interacts with NCAPG2 (NCAPD3^Tail-G2^), and the NCAPH2^Neck^ (NCAPD3^Tail-H2^), sequestering it on the surface of NCAPG2. Potential phosphosites are highlighted. Cryo-EM density is displayed over the model to illustrate local map quality. **(G)** Histograms of condensin II mass distributions in the absence of ATP (top), in the presence of 10 μM ATP (middle) and 2.5 mM ATP (bottom).

The condensin II complex forms a closed dimer mediated through contacts between the NCAPG2 subunit of one protomer and the C-terminus of the SMC4 subunit of the other (Figure 1C). We also observed modest inter-protomer interactions between NCAPH2 subunits (Figure 1D). Condensin II multimerization has been proposed to drive chromosome axis formation, and mutagenesis of this NCAPH2 region leads to poor chromosome assembly *in vitro* ^20^. As recent studies have implicated this region of NCAPG2 in binding both MCPH1 and M18BP1 ^16,17^, dimerization could also influence regulator binding.

SMC family proteins feature N and C-terminal domains that fold together to generate composite ATPase ‘head’ domains that provide motor activity upon head engagement. In the apo structure, the SMC2 and SMC4 ATPase heads are not engaged in an ATP-binding competent configuration but rather are splayed apart (Figure 1B, Figure S3A). The kleisin subunit NCAPH2 decorates the HEAT repeat subunits, and binds SMC4 through its C-terminal winged-helix domain (WHD), linking the condensin II subunits (Figure S3B). NCAPH2 residues 146-197, 314-371, and 489-593 engage NCAPD3, NCAPG2 and SMC4, respectively (Figure S3B). The region of NCAPH2 immediately emerging from the winged-helix domain SMC4 forms an additional contact with NCAPD3 within the monomer (Figure S3C). and residues 342-351 emerge from the surface of NCAPG2 to augment dimerization with SMC4-NCAPH2 of the adjacent protomer (Figure 1D).

The HEAT-repeat proteins, NCAPG2 and NCAPD3, heterodimerize using an α-helical ‘docker’ domain, important for function *in vitro* ^21^, (Figure 1A,B; Figure S3D), whereas the major SMC-HEAT contact within the protomer monomer involves binding of NCAPD3 to the surface of SMC4, including its W-loop comparable to budding yeast condensin I ^26,27^ (Figure 1B,E).

We observed map density bound to the NCAPG2 subunit that could not be accounted for by NCAPG2, and was not predicted by AlphaFold3. Automated model-building^28^ revealed that the additional density corresponds to the NCAPH2^Neck^ (Figure 1F, Figure S1C). In related eukaryotic SMCs, the kleisin^Neck^ typically binds the coiled-coil of its cognate SMC, in this case SMC2, to form Neck ‘gates’ that regulate DNA entry or exit^27,29-33^. The presence of NCAPH2^Neck^ on NCAPG2 indicates that apo condensin II possesses an ‘open’ Neck gate, which is thought to correspond to states either capturing or releasing DNA.

We observed additional density directly bridging the NCAPH2^Neck^ and NCAPG2, deriving from the C-terminal NCAPD3^Tail^, comprising amino-acids 1454-1476 (Figure 1F). NCAPD3^Tail^ residues S1458 and S1474, are buried at the NCAPH2 and NCAPG2 binding interface, respectively, and are phosphorylated in cells^34-36^, which would be incompatible with NCAPD3^Tail^ binding (Figure 1F). Consistently, deletion of the NCAPD3^Tail^ correlates with hypercondensation of chromatin *in vitro* (as examined extensively below), and alanine substitutions of the putative phosphorylation sites results in decreased activity^20^. We therefore hypothesize that NCAPH2^Neck^ sequestration on the surface of NCAPG2 by the NCAPD3^Tail^ provides a potential mechanism to regulate condensin II activity by restricting its ability to capture DNA.

### ATP-engaged condensin II is an autoinhibited monomer

Modelling ATP-dependent head engagement indicates a clash between the SMC2 and NCAPD3, which suggested dimer formation is incompatible with head engagement (Figure S3A, E). We observed that even in the presence of low micromolar concentrations of ATP, the dimer fraction of condensin II effectively dissolves in solution (Figure 1G, Figure S3F).

To understand how condensin II structurally reorganizes upon ATP-binding, a necessary aspect of its functional cycle, we determined its structure in complex with ATP to a resolution of 3.8 Å (Figure 2A, Figure S2G,H,; Figure S4 A,B). The resulting complex is monomeric, in agreement with our in-solution measurements (Figure 1G, Figure S2G). The ATPase itself adopts a conformation similar to related SMC complexes, and the presence of ATP in the active site is clearly defined in the cryo-EM map (Figure 2C,D). In this conformation, the structure of the HEAT-repeat subunit heterodimer is identical to the apo complex, however NCAPG2 replaces NCAPD3 to bind the SMC2/4 ATPase module (Figure S4C). NCAPG2 forms extensive contacts with the surface of the SMC2 ATPase domain, and additionally binds through its N-terminus to the coiled-coils of SMC4 (Figure 2B). Compared to apo condensin II, due to swapping of the HEAT repeat domains relative to the ATPase, the NCAPH2^Neck^ is repositioned in close vicinity to SMC2 (Figure 2B, Figure S4C). As a consequence, the NCAPH2^Neck^ now directly bridges NCAPG2 with the SMC2^Neck^ coils, but nevertheless remains sequestered by the NCAPD3^Tail^ (Figure 2B).

**Figure 2:**
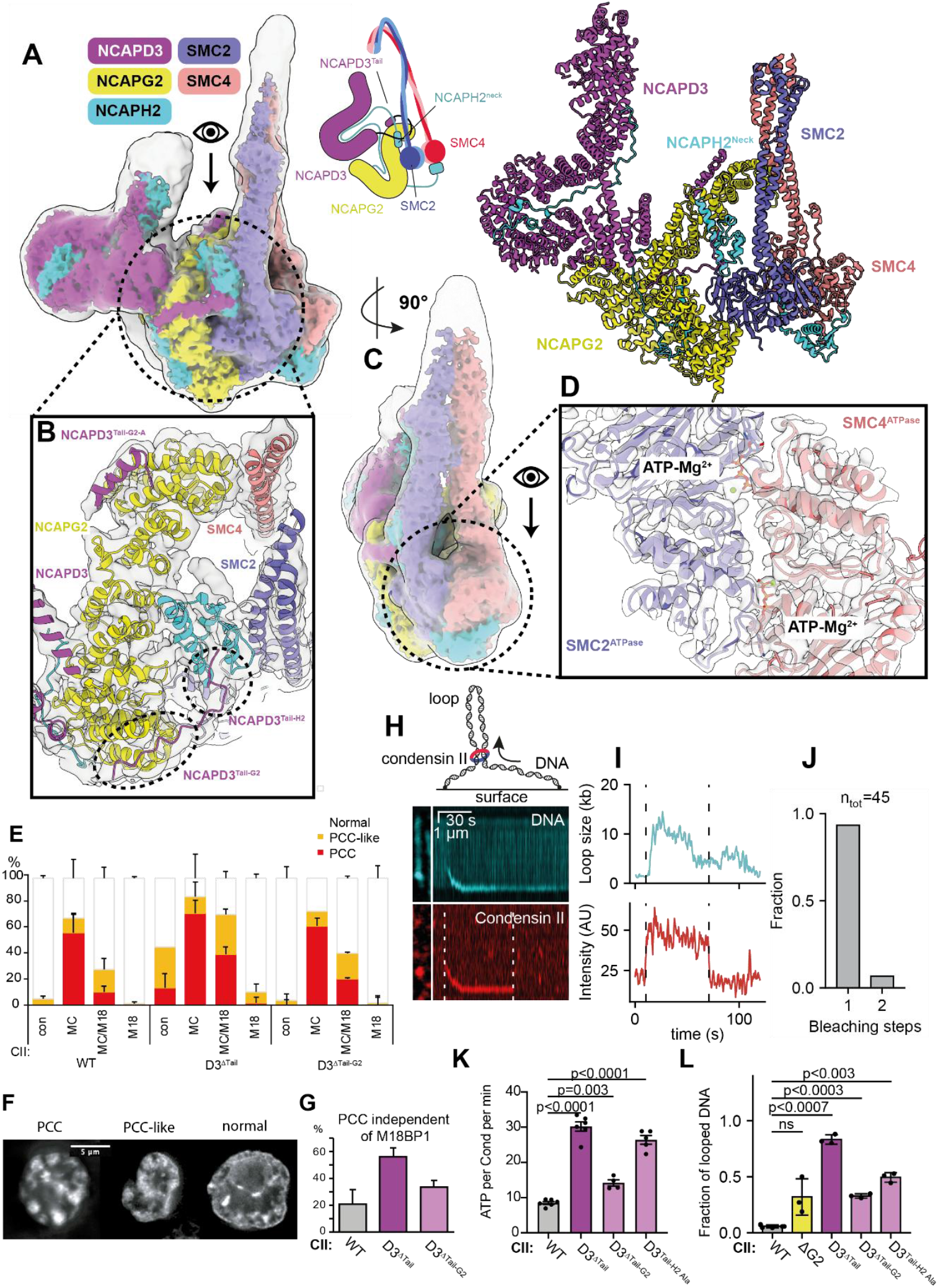
ATP bound condensin II is inhibited via interactions of the C-terminal region of NCAPD3. **(A)** Consensus cryo-EM reconstruction of condensin II bound to ATP (left panel), and corresponding molecular model (right panel). A cartoon schematic of ATP-bound condensin II is shown **(B)** Zoomed-view of cryo-EM map density, showing the trajectory of NCAPD3^Tail^. NCAPD3^Tail^ binds the N-terminus of NCAPG2 (NCAPD3^Tail-G2A^), contacts the NCAPH2^Neck^ (NCAPD3^Tail-H2^), and rebinds a second site on NCAPG2 (NCAPD3^Tail-G2^), thereby forming a ‘loop’ around NCAPH2^Neck^. **(C)** Front view of the condensin II – ATP complex. The consensus cryo-EM reconstruction is displayed as transluscent grey over a local-resolution filtered reconstruction coloured according to subunit. **(D)** Cryo-EM density contoured around the condensin II – ATP structure shows the presence of ATP and Mg^2+^ in the active site of the ATPase. **(E)** PCC assay of G2-arrested DT40 cells expressing wild-type (wt) or mutant GgCAPD3-mScarlet3. Chromosome morphology was classified as PCC, PCC-like, or normal. Guide RNAs targeted MCPH1 (M<u>C</u>), Mis18BP1/KNL2 (<u>M18</u>), or both (<u>MC/M18</u>). >50 mScarlet3-positive cells were scored per condition. Biological replicates: wt (n = 5), ΔC<u>ΔTail</u> (n = 4), 1458–1463 (n = 4). Mean + s.d. is shown. **(F)** Representative images of chromosome morphology in DT40 cells. **(G)** Ratio of PCC-positive cells in the absence of Mis18BP1/KNL2, calculated as PCC(<u>MC/M18</u>)/PCC(M<u>C</u>). The difference between wt and ΔTail was significant (Kruskal–Wallis test). **(H)** Schematic of the in vitro loop extrusion assay (top). Representative snapshots and corresponding fluorescence intensity kymographs of a condensin II-mediated loop extrusion event, showing a SxO-stained DNA molecule (middle) and Alexa 647-labeled condensin II (bottom). **(I)** Time traces of loop size (top) and condensin II fluorescence intensity (bottom) extracted from the loop extrusion event in **(H). (J)** Fraction of loop-extruding condensin II molecules exhibiting either one- or two-step photobleaching. **(K)** ATP turn-over of condensin II WT, ΔG2, D3^ΔTAIL^, D3^Δ1470-1475^ and D3^H2^ ^Alanine^ mutants, error bars indicate mean and standard error. **(L)** Fractions of DNA molecules with condensin II-mediated loops in the presence of 2 nM condensin II and 2.5 mM ATP for WT, ΔG2, D3^ΔTail^, D3^ΔTail-G2^ and D3^Tail-H2 Alanine^ mutant. Mean ± s.d. is shown. n_tot_ = 704, 292, 477, 333, 403, respectively. Data are from at least three independent experiments. The P values were calculated using Welch’s t test.

In this structure, the NCAPH2^Neck^ remains bound to NCAPG2, but a more extensive interface between NCAPG2 and NCAPD3^Tail^ are apparent. Similar to the apo structure, NCAPD3^Tail^ residues 1454-1460 (NCAPD3^Tail-H2^) bind the NCAPH2^Neck^ whereas NCAPD3^Tail^ residues 1469-1476 (NCAPD3^Tail-G2^) bind NCAPG2. However, in the ATP-bound structure, we observe the preceding residues 1336-1371 (NCAPD3^Tail-G2A^) bound to NCAPG2 (Figure 2B). By consequence, the NCAPD3^Tail^ effectively forms an extensive proteinaceous belt that encloses NCAPH2^Neck^ domain on the surface of NCAPG2 (Figure 2B).

### Condensin II is auto-inhibited *in vivo*

To establish whether the NCAPD3^Tail^ inhibits condensin II function in the cell, we implemented an assay to monitor chromosome morphology in DT40 cells arrested at the G2/M transition, prior to global chromatin compaction by condensin II, via inhibition of analogue-sensitive CDK1 ^1^. In this assay, we transiently express NCAPD3 protein (wild-type or mutants) in chicken DT40 CDK1^as^/NCAPD3-AID cells and acutely depleted endogenous chicken NCAPD3 protein (gNCAPD3) by addition of 5-Ph-IAA^37^ (Figure S4D). Cells expressing wild-type gNCAPD3 exhibited chromosome morphologies equivalent to the wild type cells during G2 and also in mitosis (Figure 2E,F). In contrast, the cells expressing gNCAPD3 with tail residues 1293-1493 deleted (gNCAPD3^ΔTail^) and residues 1458-63 deleted (gNCAPD3^ΔTail-G2^) underwent premature chromosome condensation (PCC) in G2 blocked cells (Figure 2E, F and Figure S2E), indicating premature activation of the condensin II complex.

Condensin II is repressed by MCPH1 and requires M18BP1 to condense chromatin in human cells ^16,17,25,38-40^. In order to further dissect how these factors contribute to condensin II regulation in the context of autoinhibition, we co-transfected cells with a plasmid encoding both guide RNAs (gRNAs) and Cas9 nuclease to inactivate the corresponding genes.

Following disruption of MCPH1, the majority of cells underwent PCC during G2 arrest (Figure 2F). Conversely, no PCC was observed when M18BP1 was inactivated (Figure 2F,G). Both results were consistent with their proposed roles in inhibition, and activation respectively. When both MCPH1 are depleted simultaneously, we observed marked differences between wild-type and gNCAPD3^ΔTail^ rescue conditions. Whereas MCPH1-mediated PCC was highly dependent on M18BP1 in the wild-type rescue, the gNCAPD3^ΔTail^ expressing cells exhibited PCC even in the absence of M18BP1 (Figure 2G). A more subtle mutant removing the NCAPD3^Tail-G2^ (gNCAPD3 residues 1458-1363; see Figure 2B: NCAPD3^Tail-H2^ interaction) showed an intermediate phenotype (Figure 2E). We therefore conclude that condensin II is able to spontaneously compact chromatin independently M18BP1 when the gNCAPD3^Tail^ is deleted, thereby uncovering a condensin II autoinhibition mechanism in cells (Figure 2G). A prevailing hypothesis is that condensin II complexes organize chromosomes by loop extrusion. We therefore hypothesized that the premature compaction of chromatin we observed following NCAPD3^Tail^ deletion reflects upregulation of condensin II loop-extrusion activity and sought to test this directly *in vitro*.

### Loop extrusion by condensin II is auto-inhibited by NCAPD3-tail and H2 neck interactions

To characterize the DNA-organizing activity of condensin II *in vitro*, we employed a single-molecule assay that allows direct visualization of DNA topology and condensin II complexes (Figure 2H). Upon introduction of ATP and single fluorophore labeled condensin II, we observed that wild-type condensin II bound to DNA and subsequently formed loops that gradually increased in size over time, indicative of active loop extrusion (Figure 2H,I). In the majority of cases, condensin II bound to DNA in a single step, followed by loop growth, and finally bleached in a single step (Figure 1J). This indicates that the loop-extruding form of condensin II acts as a monomer, consistent with our structure showing ATP disrupts dimerization, and prior observations^19,41^. DNA-bound condensin II complexes that did not extrude loops were also monomeric in the presence of ATP (Figure S3G), consistent with our structural findings.

To dissect whether the NCAPG2– NCAPH2^Neck^ – NCAPD3^Tail^ interaction is also autoinhibitory *in vitro*, we generated a series of mutations predicted to disrupt this interface. Specifically, we deleted the entire NCAPG2 (ΔG2) and NCAPD3 (ΔD3) subunits, separately. To more selectively perturb the NCAPD3^Tail^ we additionally removed the NCAPD3^Tail^ (NCAPD3^ΔTail^), deleted six residues that mediate the interaction with NCAPG2 (NCAPD3^ΔTail-G2^, see Fig.1F, Figure 2B: NCAPD3^Tail-G2^ interaction). Finally we introduced alanine substitutions at residues 1453-1458, inclusive, on NCAPD3 that form the H2–neck binding interface (D3^Tail-H2ala^, see Figure 1F, Figure 2B: NCAPD3^Tail-H2^ interaction). All mutants exhibited stimulated ATPase activity (Figure 2K; Figure S4F,G), compared to the wild-type.

All NCAPD3^Tail^ mutants resulted in an increase in loop extrusion activity relative to wild-type condensin II (Figure 2L), with the most prominent enhancement observed for the NCAPD3^ΔTail^.We therefore propose that the sequestration of NCAPH2^Neck^ by the NCAPD3^Tail^ autoinhibits condensin II, as ablation of this interaction promotes loop extrusion activity.

### M18BP1 activates condensin II in a CDK1-Cyclin B phosphorylation-dependent manner by releasing NCAPD3^Tail^

Recruitment of condensin II to chromosomes in mitosis involves the activator protein, M18BP1, however, it is currently unknown how M18BP1 enables condensin II activation. To understand how M18BP1 converts condensin II into an active form *in vitro*, we exploited our *in vitro* assay to determine its influence on condensin II–mediated loop extrusion. Analysis of looping probability revealed that M18BP1 alone increases loop extrusion activity only moderately, even at high molar excess (Figure 3A, Figure S5D), indicating M18BP1 by itself is not sufficient to activate condensin II.

**Figure 3:**
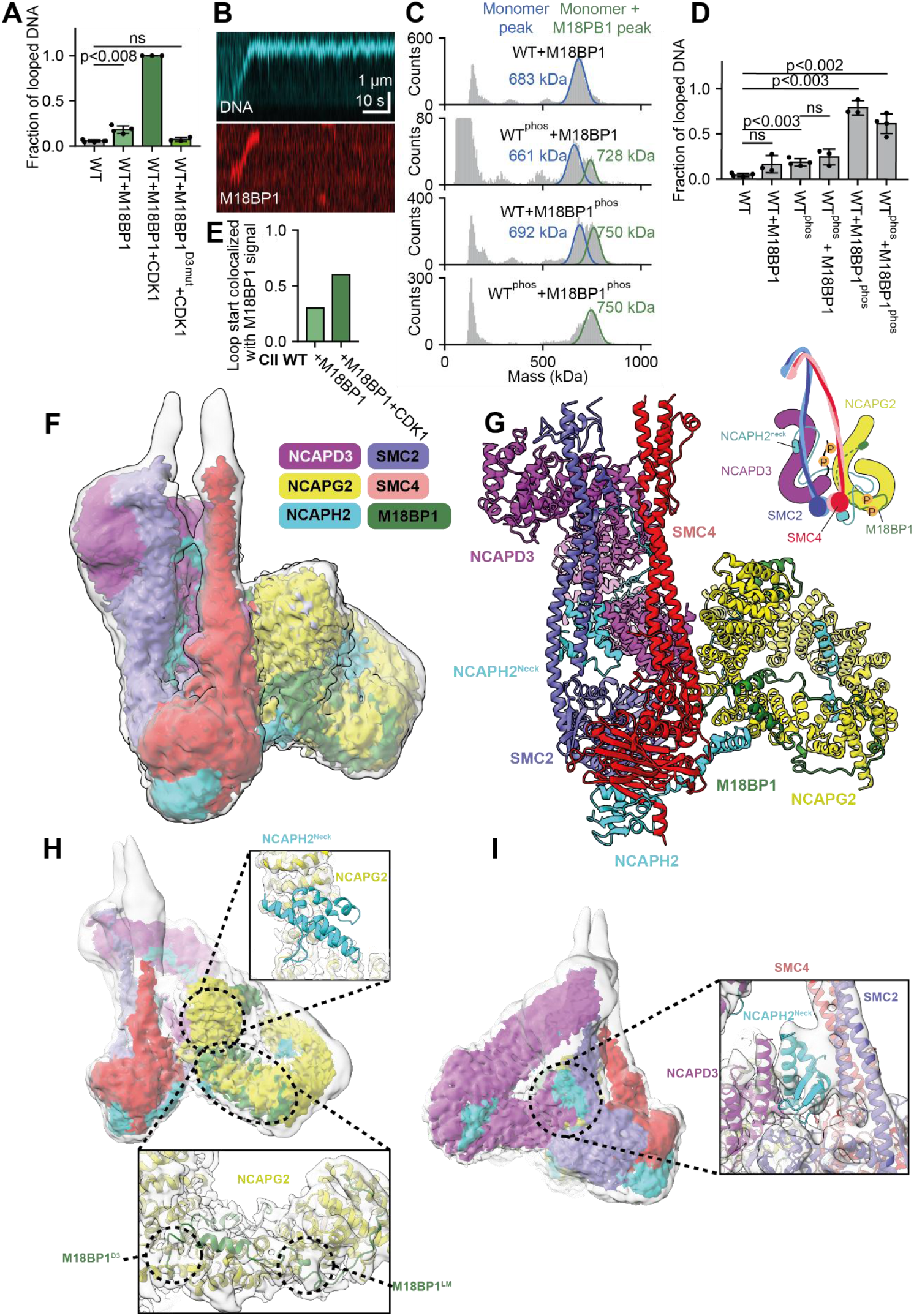
M18BP1 activates condensin II by displacing NCAPD3^Tail^, allowing condensin II DNA-clamp assembly. **(A)** Fractions of DNA molecules exhibiting condensin II-mediated loops in the presence of 2 nM condensin II and 2.5 mM ATP with either 200 nM M18BP1; 50 nM M18BP1 and 5 nM CDK1/10 nM CyclinB; or 50 nM M18BP1^D3^ ^mutant^ and 5 nM CDK1/10 nM CyclinB. Mean ± s.d. is shown. n_tot_ = 704, 523, 348, and 310, respectively. Data are from at least three independent repetitions. The data for condensin II WT are reproduced from Figure 2I. The P values were calculated using Welch’s t test. **(B)** Representative kymographs of a looping event by M18BP1 bound-condensin II in the presence of CDK1/Cyclin B, showing a SxO-stained DNA molecule (top) and Janelia Fluor 646-labeled M18BP1 (bottom). **(C**) Histograms of mass distributions for the indicated samples: WT+M18BP1, WT_phos_+M18BP1, WT+M18BP1_phos_ and WT_phos_+M18BP1_phos_. Representative datasets from multiple independent experiments are shown. **(D**) Fractions of DNA molecules exhibiting condensin II-mediated loops in the presence of 2 nM condensin II and 2.5 mM ATP for WT, WT+M18BP1, WT_phos_, WT_phos_+M18BP1, WT+M18BP1_phos_ and WT_phos_+M18BP1_phos_. Mean ± s.d. are shown. n_tot_ = 774, 533, 798, 374, 564, 754 respectively. Data are from at least three independent experiments. The P values were calculated using Welch’s t test. **(E)** Normalized fraction of condensin II WT mediated loops, colocalized with M18BP1 fluorescence signal in the presence of 2 nM condensin II WT and 10 nM Janelia Fluor 646 labelled M18BP1, in the absence and presence of 5 nM CDK1 and 10 nM Cyclin B. n_loop_ = 20 and 35, respectively. **(F)** Consensus cryo-EM reconstruction of condensin II bound to ATP and M18BP1, and corresponding molecular model **(G)**. A cartoon schematic of ATP/M18BP1-bound condensin II is shown. (P) signifies phosphorylation. **(H)** Structure of CII bound to M18BP1 and ATP. Upper right inset shows cryo-EM map density contoured around NCAPG2. In the activated state, there is no map density for NCAPH2^Tail^ on NCAPG2, indicating it has been released. Lower right panel: cryo-EM density for M18BP1 indicates it binds multiple sites on NCAPG2, including a site overlapping with where NCAPD3^Tail^ binds in the inhibited state. **(I)** Rotated view of the M18MP1/ATP-bound CII structure. Inset panel shows cryo-EM map density contoured around NCAPH2^Neck^ which has relocated to the SMC2^Neck^ region.

CDK1/Cyclin B phosphorylation is important for condensin II activation *in vivo*^22,23^, hence we reasoned that it may also be fundamental to DNA looping activity *in vitro*. When condensin II and M18BP1 were pre-incubated with CDK1/Cyclin B prior to imaging, we observed a marked stimulation of looping activity, with nearly all DNA molecules being organized into loops (Figure 3A). Control experiments with Janelia Fluor 646-labeled M18BP1 confirmed that loop formation resulted from specific association of M18BP1 with condensin II (Figure S6). Together, these results indicate that CDK1/Cyclin-B substantially augments the capacity of M18BP1 to activate condensin II.

To directly examine the role of phosphorylation, we analyzed samples in which either condensin II or M18BP1 was individually or both phosphorylated (Figure 3C,D). Mass photometry showed that phosphorylation of either component increased binding affinity, with the strongest effect observed upon phosphorylation of M18BP1 (Figure 3C). The subsequent looping probability analysis revealed that phosphorylation of M18BP1 alone is sufficient to stimulate loop extrusion to a level comparable to the doubly phosphorylated sample (Figure 3D). Consistent with this result, single-molecule imaging revealed increased localization of M18BP1 at loop sites following CDK1/Cyclin B treatment, indicating enhanced binding affinity^16^ (Figure 3B,E; Figure S6A). Mass spectrometry revealed that phosphorylation sites on M18BP1 are located near positively charged regions of NCAPG2, providing a structural explanation for how phosphorylation enhances binding (Figure S7A).

Having established conditions for efficient in vitro activation of condensin II by M18BP1, we next asked how M18BP1 association leads to loop extrusion activation. Overlaying AlphaFold3^42^ predictions of M18BP1 and NCAPG2 with our experimental structure of auto-inhibited condensin II shows clear overlap between the NCAPD3^Tail^ residues 1470-1475 and amino acids 1073-1078 of M18BP1 (Figure S5A-C), suggesting that M18BP1 may activate condensin II by displacing the NCAPD3^Tail^, and, consequently the NCAPH2^Neck^. Consistently, we observed truncation of NCAPD3 tail residues 1470–1475 results in increased loop-extrusion activity (Figure 2L) and PCC *in vivo* (Figure 2E,G).

Supporting this hypothesis, hyperactivation was completely eliminated by point mutations that specifically target the ability of M18BP1 to bind the NCAPD3^Tail^ overlapping region on NCAPG2 (M18BP1^D3^ ^mutant^), without impacting M18BP1 recruitment to condensin II (Figure 3A, Figure S5A-G). In addition, mass spectrometry detected more NCAPD3^Tail^ phospho-peptides, including key interfacial residues S1458 and S1474 (consistent with our competition model suggesting M18BP1 binding drives NCAPD3^Tail^ release, which would presumably render it accessible to phosphorylation.

Together, these data indicate that M18BP1 phosphorylation plays the major role in activating condensin II. It does so by enhancing its binding affinity to the complex, enabling it to displace the autoinhibitory NCAPD3^Tail^, in turn increasing the susceptibility of the latter to phosphorylation. We hypothesized that NCAPD3^Tail^ release would in turn liberate the NCAPH2^Neck^ to activate condensin II, and therefore applied cryo-EM to establish the molecular basis by which M18BP1 achieves condensin II activation.

### M18BP1 releases the NCAPH2^Neck^ and reorganizes condensin II into a state ‘poised’ for DNA capture

To understand the structural mechanism by which M18BP1 activates condensin II, we assembled condensin II with M18BP1, phosphorylated the complex with CDK1/cyclin B, and determined its structure in the presence of ATP (Figure 3F, G, Figure S7C, D). We observed both ATP-engaged and non-engaged forms of the SMC2/4 ATPases, to a resolution of 4.4 Å and 4.5 Å (Figure S2I-N), respectively. Consistent with our hypothesis and single-molecule experiments, NCAPD3^Tail^ is displaced by M18BP1, this is driven by competition by M18BP1, which now occupies the NCAPD3^Tail^ binding site (Figure 3H, Figure S5A,E,) and forms a highly extended interface with NCAPG2 (Figure 3H, lower inset). In agreement with the notion that NCAPD3^Tail^ autoinhibits via sequestration, in the presence of M18BP1, NCAPH2^Neck^ has clearly also been released from NCAPG2 (Figure 3H, upper inset).

The released NCAPH2^Neck^, is now wedged between the C-terminal alpha-helices of NCAPD3 (Figure 3I) and remains bound to the coiled-coils of the SMC2, in a conformation similar to the auto-inhibited state. The NCAPD3^Tail^ beta-strand that interacts with NCAPH2^Neck^ in the auto-inhibited state was absent (Figure 3I), potentially as a result of incompatible phosphorylation of NCAPD3^Tail^ residue S1458 (Figure 1F, Figure S7B).

Following activation by M18BP1, the reorganization of the HEATs relative to the ATPase domains (Figure S7E) positions the NCAPD3 in close proximity to the coiled-coils of SMC2^Neck^. In related eukaryotic SMCs, domains equivalent to SMC2^Neck^:NCAPH2^Neck^ gate cooperates with the cognate HEAT, in this case NCAPD3, to form an ATP-dependent DNA-binding clamp^29,43-45^. Mapping of surface electrostatics showed that the NCAPD3 presents a positively charged surface to solvent in close proximity to the SMC2^Neck^:NCAPH2^Neck^ gate (Figure S7F). M18BP1 thereby appears to facilitate reorganization of condensin II into a conformation that is poised to capture DNA. We therefore anticipated that condensin II might interface with DNA by assembling a structurally analogous ‘clamp’.

### Condensin II forms a DNA-binding clamp

To understand how condensin II reorganizes to bind DNA, we initially attempted to obtain a structure of condensin II bound to M18BP1 in the presence of DNA and phosphorylation, however the number of intact particles was limiting (Figure S8A,B), resulting in a reconstruction limited to 15 Å. By contrast, complexes of CDK1/CyclinB phosphorylated condensin II assembled together with DNA and the transition state nucleotide analogue ADP.BeFx yielded a cryo-EM reconstruction of the DNA-bound state to an overall resolution of 3.8 Å (Figure S2O,P;Figure S8C,D; Figure 4A,B).

**Figure 4:**
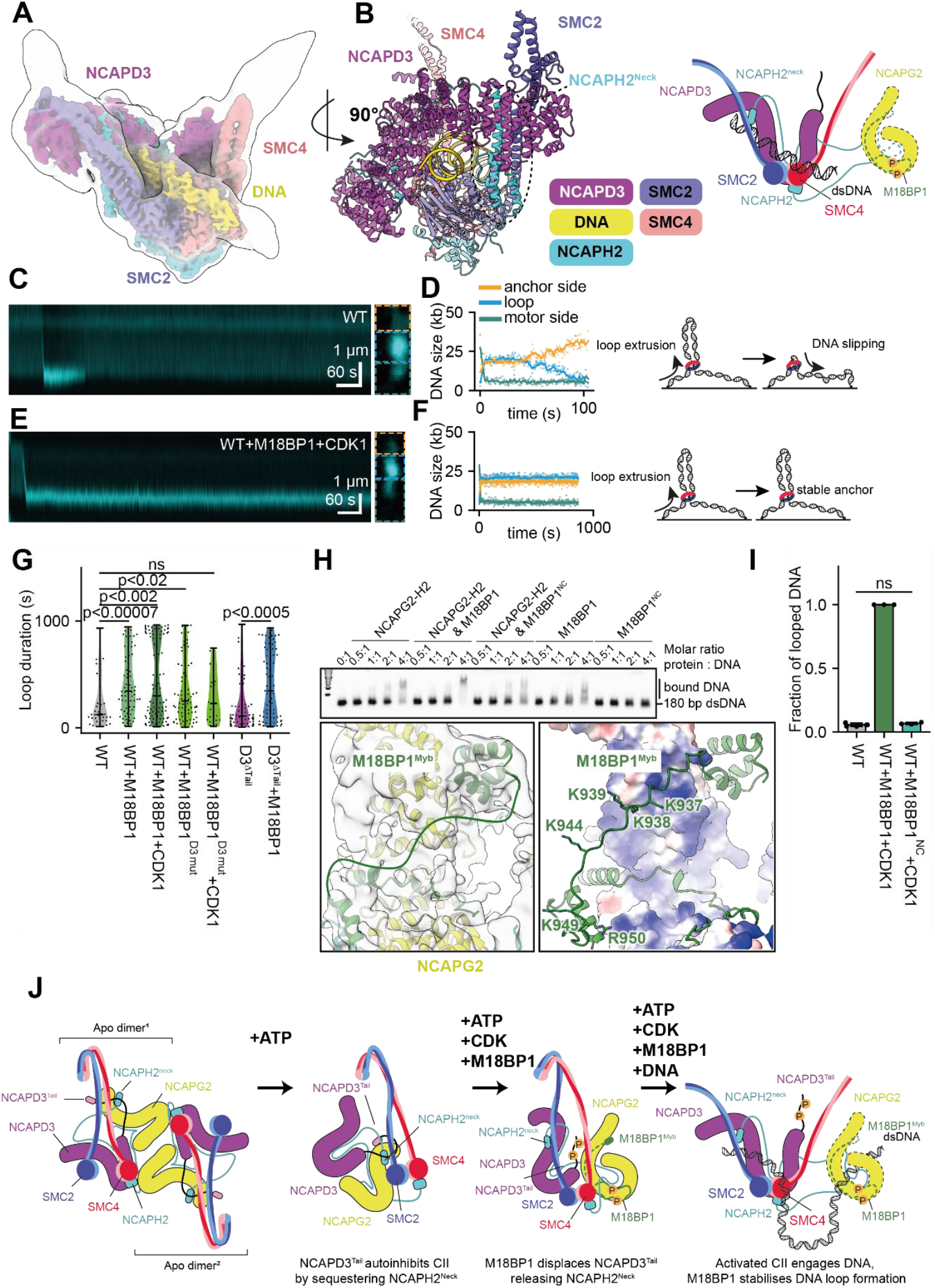
M18BP1 stabilizes condensin II loops by providing a DNA anchor. **(A)** Consensus cryo-EM reconstruction of the condensin II – ADP.BeFx – DNA complex. A local-resolution filtered reconstruction is colored according to subunit assignment. The SMC2/4 coiled coils are splayed apart compared to other structural states, and DNA is bound across a composite surface formed by the condensin II ATPase domains. A cartoon schematic is shown (P) signifies phosphorylation **(B)** Rotated view of local-resolution filtered map density for the condensin II – ADP.BeFx – DNA complex coloured according to subunit. DNA is topologically enclosed by formation of a clamp. The compartment is sealed from above by the NCAPD3 subunit that forms a tripartite DNA-binding interface with the NCAPH2^Neck^:SMC2^Neck^, and below by the SMC2/SMC4 ATPase domains. **(C, E)** Representative kymograph of a loop extrusion event by WT condensin II (C) and by condensin II in the presence of M18BP1 and Cyclin B/CDK1 (E), showing SxO-stained DNA molecules. **(D, F)** DNA lengths extracted from the corresponding kymographs in (C) and (E), respectively, for regions outside the loop (motor and anchor sides) and within the loop (Loop). **(G)** Loop duration times for loops formed by WT condensin II and in the presence of M18BP1; M18BP1 plus CDK1, M18BP1^D3^ ^mutant^ and M18BP1^D3^ ^mutant^ plus CDK1. n=41, 78, 106, 86, 29, respectively. The P values were calculated using Wilcoxon rank-sum test. **(H)** Electrophoretic mobility shift assay (EMSA) of an NCAPG2-NCAPH2 subcomplex in the presence or absence of indicated M18BP1 variants titrated against DNA (upper panel). Molar ratios and the position of unbound vs. shifted DNA are indicated. Data are from at least three independent replicates. A zoomed view of the NCAPG2-NCAPH2-M18BP1 subcomplex in the ATP/M18BP1-condensin II structure with cryo-EM map density displayed translucently shows separation of two M18BP1 regions connected by a flexible linker, drawn as a cartoon (lower left). Surface charge of NCAPG2 and positions of positively charged residues within an AlphaFold3^42^ prediction of M18BP1 are displayed (lower right) **(I)** Fractions of DNA molecules exhibiting condensin II-mediated loops in the presence of 2 nM condensin II and 2.5 mM ATP with either 50 nM M18BP1 and 5 nM CDK1/10 nM CyclinB or 50 nM M18BP1^NC^ ^mutant^ and 5 nM CDK1/10 nM CyclinB. Mean ± s.d. is shown. n_tot_ = 704, 348, and 390, respectively. Data are from at least three independent repetitions. The data for condensin II WT and condensin II WT + M18BP1 + CDK1 are are reproduced from Figure 3A. The P values were calculated using Welch’s t test. **(J)** Cartoon schematic of condensin II activation. (P) signifies phosphorylation. Apo condensin II reorganizes into a monomer upon binding of ATP. Sequestration of NCAPH2^Neck^ by NCAPD3^Tail^ maintains auto-inhibition. Binding of M18BP1 to condensin II displaces NCAPD3^Tail^, exposing it for phosphorylation. Displacement of the NCAPD3^Tail^ releases NCAPH2^Neck^, which is then able to participate in DNA clamp formation, and downstream DNA organization. DNA binding by the NCAPG2-NCAPH2-M18BP1 subcomplex stabilizes loop formation.

In this structure, DNA is ‘clamped’ between a composite interface formed by the ATP-engaged SMC2/4 heads, the NCAPH2^Neck^ and Neck coiled-coils of SMC2, and sealed above by NCAPD3 (Figure 4A,B), forming an enclosed DNA chamber. The NCAPH2^Neck^ residues 53-85 undergoes dramatic refolding in the presence of DNA, to form a triple α-helical bundle with the SMC2^Neck-coils^. This extended α-helix from NCAPH2^Neck^ results from refolding of the kleisin around a fulcrum centered around Tyr67 (Figure 4B, Figure S8E). Interestingly, in the compact NCAPH2^Neck^ structures (apo, ATP-engaged, activated complexes), the N-terminal ∼60 residues preceding this fulcrum fold to structurally mimic the SMC2^Neck-Coils^ (Figure S8F).

NCAPD3 forms extensive contacts with the SMC2-NCAPH2 neck, and also directly bridges the coiled-coils of SMC2 and SMC4 emerging from the ATPases, which are now splayed apart (Figure 4B). In this reconstruction, the NCAPG2 subunit is no longer visible. Instead, we observe that the NCAPD3-NCAPG2 dimerization interface is disrupted by binding of NCAPD3 and the SMC4 coiled-coils, indicating that HEAT hetero-dimerization is incompatible with clamped state assembly (Figure S8G).

Comparison of the DNA-clamped state with the autoinhibited structure reveals further features of autorepression, in which NCAPG2-NCAPH2^Tail^ effectively block DNA engagement via two contacts. Firstly, NCAPG2-NCAPH2^Tail^ occupy the binding site on SMC2 through which NCAPD3 docks to bind DNA (compare Figure S7F, and Figure S10 D, left). Secondly, binding of NCAPG2 to SMC2/4 additionally blocks the DNA binding site on the ATPase domains (Figure S10E, right). Because NCAPH^Tail^ bridges NCAPG2 and SMC2, its release presumably enables the conformational reorganization of the HEAT repeat subunits required for DNA capture.

A lower resolution 3D reconstruction, of the condensin II – M18BP1 – DNA complex (compare Figure S8B, D) suggests that the overall organization of the central condensin II DNA clamp is structurally similar in the presence or absence of M18BP1 (Figure S8A-D). This is consistent with our observation that NCAPG2, which is essential for M18BP1 binding to condensin II (Figure S6A-C), is disengaged from NCAPD3 in the clamped state (Figure 4A,B). The primary structural role of M18BP1 in directing formation of the active clamped state therefore appears to be NCAPH2^Neck^ release.

### M18BP1 stabilizes condensin II–mediated loops by augmenting an NCAPG2-DNA anchor

To determine whether condensin II regulatory mechanisms influence loop dynamics, we analyzed loop kinetics for condensin II variants tested in this study in single-molecule assays. We observed that the rate of loop extrusion was not significantly different across different condensin II variants (Figure S9A). In contrast, we observed significant differences in loop stability depending on the presence or absence of M18BP1 (Figure 4C-G).

Wild-type condensin II functioned as a unidirectional loop extruder, in which DNA is reeled in from one side of the complex (termed the motor side, Figure 4D). DNA on the opposite, non-extruding side of the complex (termed the anchor side, Figure 4D), however, frequently slipped out of the extruded loops (Figure 4C,D, Figure S9B), resulting in rapid loop loss (Figure 4C,D, Figure S9B) and short loop lifetimes (Figure 4G). These results indicate that, while condensin II can form loops in the absence of M18BP1 (albeit with a considerably lower probability), these loops are fundamentally unstable.

In the presence of M18BP1 and CDK1, condensin II formed markedly more stable loops (Figure 4E,F, Figure S9C), characterized by increased loop lifetimes (Figure 4G). In the presence of M18BP1, the DNA length remained constant at the anchor side over far more extended periods compared to condensin II alone (Figure 4F, Figure S9C). indicating that stabilization was primarily provided by improved anchoring of DNA.

To explicitly disentangle the role of M18BP1 in activation from that in increasing loop stability, we performed two separation of function experiments. Firstly, we tested addition of the M18BP1^D3^ ^mutant^, which still binds condensin II but does not increase looping probability (e.g. activation). Secondly, we tested whether addition of M18BP1 to complexes deleted for the autoinhibitory NCAPD3^Tail^, which are constitutively derepressed, alters loop stability. In both scenarios, we observed M18BP1-mediated loop stabilization (Figure 4G). This indicates the effect of M18BP1 in augmenting loop lifetime, is apparently distinct from its role in NCAPD3^Tail^ displacement, hence M18BP1 does not simply relieve autoinhibition, but also contributes directly to DNA loop anchoring.

In our structure of M18BP1 bound condensin II, we observed additional globular density bound to the NCAPG2 N-terminus (Figure 4H, lower left inset), that originates from the M18BP1^Myb^ domain (Figure 4H, lower left inset). Connectivity of M18BP1^Myb^ to the remaining C-terminal regions of M18BP1 binding NCAPG2 is achieved by a ∼40 amino-acid long linker that is rich in basic residues, in effect generating a positively charged loop (Figure S10A-C). As this linker passes over a highly positively charged-patch on NCAPG2 (Figure 4H, lower right inset), we therefore hypothesized the M18BP1–NCAPG2 subcomplex may act as a composite DNA-binding anchor in condensin II. To test this hypothesis, we first performed electrophoretic mobility shift assays (EMSAs) to establish the DNA-binding properties of an NCAPG2-NCAPH2 subcomplex, and the modulation thereof by M18BP1. We observed that NCAPG2-H2 and M18BP1 individually shift DNA only weakly, indicative of low binding affinity (Figure 4H, upper panel). In contrast, the NCAPG2-H2-M18BP1 trimer robustly shifts DNA, showing it indeed forms a composite DNA-binding site.

To test the role of the basic linker region in M18BP1, we inserted charge-reversal mutations into linker residues K937E, K938E, K939E, K944E, K949E, R950E (M18BP1^NC^;Figure S10B,C). The charge-reversal mutations impaired DNA binding by M18BP1 alone and reduced its ability to augment DNA binding by the NCAPG2–NCAPH2 subcomplex (Figure 4H, upper panel), without impacting binding of M18BP1 to condensin II (Figure S5F,G). Subsequently, analysis of looping probability revealed that the M18BP1^NC^ mutations caused a significant reduction in the M18BP1-mediated activation of condensin II loop extrusion to the level of wild-type condensin II (Figure 4I).

Together, these results reveal an unexpected bipartite role for M18BP1 in both overcoming autoinhibition, and generating a composite DNA-anchor together with NCAPG2-NCAPH2 that substantially stabilizes condensin II loop extrusion activity.

## Discussion

In this study, we define a mechanistic framework for the auto-inhibition and activation of human condensin II (visually summarised in Figure 4F), representing the first complete description of a regulatory pathway for any eukaryotic SMC complex. We show that condensin II is held in an inactive state through a mutually repressive configuration centered on the NCAPD3^Tail^ which sequesters the NCAPH2^Neck^ and prevents formation of an ATP-dependent DNA clamp.

Activation proceeds through a two-step mechanism in which CDK1-dependent phosphorylation of M18BP1 enables binding of the activator M18BP1, leading to displacement of the NCAPD3^Tail^, liberation of the NCAPH2^Neck^, and reorganization of condensin II into a DNA-clamping–competent conformation. Beyond ablating auto-inhibition, M18BP1 unexpectedly mechanically stabilizes loop extrusion. Together, these findings reveal how condensin II activity is both switched on and stabilized conferring novel biochemical activity specifically during mitosis.

In addition, our study provides structural evidence of human SMC complex dimerization. The condensin II dimer is not a loop extruding complex, as has been observed for other eukaryotic SMCs in single-molecule assays ^46,47^ and for cohesin in cells ^48^. Recent evidence suggests that segments of NCAPH2 at the dimer interface identified here are important for chromosome axis assembly ^20^. Although speculative, dimerization could therefore have an architectural role in guiding chromosome formation. Furthermore, as one dimerization contact involves SMC4, common to both condensin I and II, this introduces the possibility that both condensins could physically interact to coordinate chromosome structure.

Condensin II is constitutively nuclear, yet only stably binds and globally compacts DNA during mitosis^1,2,14,17,49,50^, although it may have a role in establishing or maintaining interphase chromosome territories^51,52^. Premature activation of condensin II causes aberrant genome structure and pathology, including microcephaly^17,38-40,53-55^. How condensin II is regulated to prevent premature formation of mitotic chromosomes has therefore remained an outstanding mechanistic question.

Activation of condensin II is tightly coupled to mitotic entry through phosphorylation ^22,23^, primarily by CDK1, but also Mps1 ^24^,.

We propose CDK1 phosphorylation has a dual role in activation. Firstly, it enhances the capacity of M18BP1 to activate condensin II by releasing the NCAPH2^Tail^, via NCAPD3^Tail^ displacement. Secondly, CDK1 phosphorylation of NCAPD3^Tail^ could prevent NCAPD3 from rebinding NCAPH2^Neck^ and NCAPG2, to delay reestablishment of autoinhibition.

Across eukaryotic SMCs, NCAPG2 orthologues, or paralogous equivalents (broadly termed ‘HAWK-B’ subunits), are a focus of regulator recruitment ^16,56-61^ and can bind DNA independently ^44,62-66^. Regulation of chromatin association of condensins^16,56,57^ and cohesins^58-61^ via HAWK-B thus is emerging as a general mechanism for control of genome folding by eukaryotic SMCs. Our study suggests autoinhibition and activation of human condensin complexes have similar mechanistic features. In condensin I, the N-terminus of the kleisin is bound to NCAPG^57^ and loss of this region results in hyperactivity^67^. Similarly, the C-terminal tail of NCAPD2 contributes to autoinhibition and competes with binding of activator KIF4A^57^. As in condensin II, these interactions are primarily driven by extended disordered regions, which harbor multiple cell cycle specific phosphorylation sites ^57^.

The budding yeast condensin I HAWK-B subcomplex features a ‘safety-belt’ in which the kleisin forms a proteinaceous loop anchoring condensin to DNA ^62,64^. In contrast, we found human condensin II appears to lack a kleisin-mediated safety belt. Instead our results suggest that the M18BP1 provides an extrinsic, regulatable safety-belt to augment DNA-binding and stabilize loop formation. Delegation of DNA loop anchoring to the extrinsic factor M18BP1, whose association with the complex can be regulated via several mechanisms, affords exquisite control over condensin II activation. As condensin I/II and cohesin utilize comparable HEAT-repeat hubs to recruit regulators across eukaryotes, it will be of fundamental interest to determine whether similar extrinsic mechanisms underly all aspects of SMC-lead genome organization.

## Data and Materials Availability

All imaging data, cell lines and plasmids are available upon request to the corresponding authors. Cryo-electron microscopy density maps were deposited with the Electron Microscopy Data Bank (EMDB). Corresponding atomic coordinates were deposited with the Protein Data Bank (PDB).

## Acknowledgements

This work was funded by the Wellcome Trust (311085/Z/24/Z to KWM, and 107022/Z/15/Z to WCE), Sheffield University Physics of Life Fellowship (EEC), Medical Research Council UKRI MC-A652-5PY00 (LA) and MC_UP_1201/6 (DB), the Max Planck Society (EK), European Research Council Starting Grant 101076914 (EK) Deutsche Forschungsgemeinschaft (DFG, German Research Foundation)—SFB 1551 (DT and EK), and Cancer Research UK C576/A25675 (DB).

We acknowledge Diamond Light Source for access and support of the cryo-EM facilities at the UK’s national Electron Bio-imaging Centre (eBIC) [under proposal EM BI31336], funded by the Wellcome Trust, MRC and BBRSC. This work was supported by funding for the Wellcome Discovery Research Platform for Hidden Cell Biology [226791] and we gratefully acknowledge support from the Proteomics and Structural Biology Cores. We thank the Vannini group (Human Technopole) for communicating unpublished results. We thank members of the Edinburgh chromosome supergroup for discussions. We thank Mohit Misra and Ivan Đikić for providing access to the mass photometry instrument. We thank Gabriele Maul for the preparation of biotinylated DNA.

## Author Contributions

Conceptualization: E.C, E.K, K.W.M, K.S., D.T., Methodology: E.C, E.K, Y.L., K.W.M, K.S., D.T., Investigation: E.C, E.K, Y.L., K.W.M, D.S., K.S., D.T., Visualization: E.C,E.K, Y.L.,K.W.M, K.S.,D.T., Funding acquisition: L.A.,D.B.,E.C, W.C.E, E.K, K.W.M., Project administration: E.C., E.K., K.W.M., Writing – original draft: E.C, E.K, K.W.M, K.S., D.T., Writing – review & editing: E.C, W.C.E., E.K, K.W.M, Y. L., K.S., D.T.

## Declaration of Interests

The authors declare no competing interests.

## Notes

### Competing Interest Statement

The authors have declared no competing interest.

### Summary of Updates

Main and supplemental text updated for clarity. Single molecule and structural data presented in figs 1,2, and supplement 3 reorganised.

