## Supplemental materials for "A bipartite mechanism for condensin II activation in mitosis"

#### Materials and Methods

AlphaFold modelling

AlphaFold prediction shown in Figure S5 was attained using AlphaFold3[68] using NCAPG2 residues 1-1143, NCAPH2 residues 312-420 and M18BP1 residues 841-1132. Extended regions of low confidence were removed from view for figure clarity.

Protein complex purification for cryo-electron microscopy (cryo-EM)

Genes encoding human condensin II subunits were cloned into a modified pF1 vector with a TEV-cleavable 2xStrepII tag inserted in frame with the NCAPH2 C-terminus[69].
Cells expressing condensin II and associated mutants were lysed with a sonicator in lysis buffer (40 mM HEPES pH 7.8, 1 mM EDTA, 1 mM EGTA, 300 mM NaCl, 0.5 mM TCEP and 5% glycerol) supplemented with benzamidine, EDTA-free protease inhibitor tablets (Roche) and benzonase (Sigma). Lysate was clarified by centrifugation at 38k R.C.F. in a JLA 16.25 rotor, and the resulting supernatant was loaded onto a StrepTactin column (Qiagen). The resin was then washed with 20 volumes of lysis buffer, and the complexes were eluted in lysis buffer supplemented with 5 mM desthiobiotin (IDT).

For further purification, the StrepTactin eluate was then diluted 1:1 in heparin buffer (40 mM Tris pH 7.8, 1 mM EDTA, 1 mM DTT, 5 % Glycerol) and applied to a HiTrap Heparin HP (Cytiva) column equilibrated with heparin buffer containing 150 mM NaCl. Immobilized proteins were then washed with 20 column volumes of heparin buffer plus 250 mM NaCl, followed by step elution in heparin buffer plus 500 mM NaCl.

Fractions containing condensin II were then harvested and concentrated to a volume of 5 mL, then applied to a Superose 6 16/60 (Cytiva) column pre-equilibrated in size-exclusion buffer (150 mM NaCl, 20 mM HEPES pH 7.8, 5% Glycerol, 1 mM TCEP). Relevant fractions were then pooled, concentrated, flash-frozen in liquid nitrogen and stored at -70 C until required.

Protein complex stabilization for cryo-EM

On-column crosslinking:
To stabilize the apo condensin II and DNA-clamping state, we performed on-column crosslinking. Briefly, a Superose 6 increase 3/150 GL column (Cytiva) coupled to a MicroAkta (Cytiva) was preequilibrated in crosslinking buffer (80 mM NaCl, 20 mM HEPES pH 7.8, 1 mM TCEP, 2 mM $MgCl_2$). Prior to application to the gel filtration column, the condensin II DNA-clamping complex was incubated together with a 171bp α-satellite DNA [70], and 0.1 µM CDK1:CyclinB:Cks1 in gradient buffer and phosphorylated at 30 °C for 2 hours in an Eppendorf Thermomixer, followed by addition of ADP.BeFx and a further 30 minute incubation.

100 µL 2.5 mM BS(PEG)$_5$ was injected into the MicroAkta system for 0.6 mL at a flow rate of 0.025 µL/min. Each condensin II complex was then injected into the system and eluted at the same flow rate into a 96-well plate. Each plate well was supplemented with 5 µL 1M tris pH 7.4 to quench the crosslinker during elution. Fractions containing condensin II were pooled, buffer exchanged into crosslinking buffer and concentrated (Amicon 100kDa, 0.5 mL). Following concentration, condensin II complexes were again centrifuged at 10 °C in a table-top centrifuge for 10 min, and then immediately vitrified for structural analyses.

Glycerol density gradient fixation:
For the ATP-engaged autoinhibited, and the ATP-engaged activated M18-bound condensin II states, complexes were stabilized using identically prepared glycerol density gradients prior to vitrification on cryo-EM grids.

In brief, 2 ml gradients were prepared in a buffer of 150 mM NaOac, 40 mM HEPES pH 7.5, 2 mM MgOac, and 0.5 mM TCEP, with 10% glycerol in the light solution, and 30% glycerol plus 0.08% glutaraldehyde in the heavy solution. For ATP-engaged autoinhibited CII, the gradients were additionally supplemented with 0.5 mM ATP.

Following sequential addition of the light, then heavy solution into Beckman 2.2 ml tubes, the gradients were mixed using a BioComp Gradient Master. The gradients were then allowed to rest at 4 °C for 30 minutes prior to launching ultracentrifugation.

For ATP-engaged autoinhibited condensin II, protein complex was mixed 1:1 with gradient buffer and incubated on ice for 30 minutes prior to application to the top of the glycerol gradient. For the ATP-engaged activated M18-bound condensin II complex, M18 and condensin II were incubated together with 0.1 µM CDK1:CyclinB:Cks1 in gradient buffer and phosphorylated at 30 °C for 2 hours in an Eppendorf Thermomixer, prior to application to the glycerol gradient. Both gradients were run in a Beckman Optima Benchtop Ultracentrifuge using a TLS-55 rotor, at 55k RPM, and a temperature of 4 °C for 3 hours. Fractions were harvested manually into a 96-well plate, with each plate well supplemented with 5 µL 1M tris pH 7.4 to quench the crosslinker. Fractions containing condensin II complexes were pooled, buffer exchanged into gradient buffer and concentrated (Amicon 100kDa, 0.5 mL) to a final volume of ~40 µL. Following concentration, condensin II complexes were again centrifuged at 10 °C in a table-top centrifuge for 10 min, and then immediately vitrified for structural analyses.

Cryo-electron microscopy grid vitrification

Apo-condensin II, condensin II DNA-clamping state

3 µl of prepared complexes at a concentration of approximately 1 mg/ml was applied to Quantifoil 300 mesh copper R1.2/1.3 grids (Quantifoil Micro Tools), glow discharged with an Edwards S150B glow discharger for 1 min 15 sec, setting 6, 30-35 mA, 1.2 kV, 0.2 mBar (0.15 Torr). The grids were then flash frozen in liquid ethane using a ThermoFisherScientific Vitrobot IV (2.5 s blotting time, -7 blotting force) using Whatman filter paper 1.

ATP-engaged condensin II, ATP-engaged M18BP1-bound condensin II

3 µl of prepared complexes was applied to Quantifoil 300 mesh copper R1.2/1.3 grids (Quantifoil Micro Tools), glow discharged with a PELCO easiGlow (Ted Pella) glow discharger for 2 minutes, 28 mA, 1.2 kV, 0.2 mBar (0.15 Torr). The grids were then flash frozen in liquid ethane using a ThermoFisherScientific Vitrobot IV (3.5 s blotting time, 4 blotting force) using Whatman filter paper 1.

Cryo-electron microscopy data collection

For all complexes micrograph movies were collected using EPU (ThermoFisherScientific) with AFIS activated. Defocus values ranged from -0.8 to -2.2 µm, at an interval of 0.2 µm.

Apo condensin II:

46,321 micrograph movies were collected on eBIC Titan Krios III coupled to a Falcon IV detector and Selectris X energy filter, with an operating acceleration voltage of 300 keV, a nominal magnification of 130k and pixel size of 0.931 Å, at a dose rate of 12 $e^{-}$/Å/s for a total overall dose of 40 $e^{-}$/Å$^2$.

Engaged inhibited state:
19,758 micrograph movies were collected on a Glacios 2 coupled to a Falcon IV detector and Selectris X energy filter, with an operating acceleration voltage of 200 keV, a nominal magnification of 165k and pixel size of 0.69 Å, at a dose rate of 13 $e^{-}$/Å/s for a total overall dose of 50 $e^{-}$/Å$^2$.

ATP-engaged activated M18BP1-bound condensin II:
71,128 micrograph movies were collected on a Glacios 2 coupled to a Falcon IV detector and Selectris X energy filter, with an operating acceleration voltage of 200 keV, a nominal magnification of 165k and pixel size of 0.69 Å, at a dose rate of 13 $e^{-}$/Å/s for a total overall dose of 50 $e^{-}$/Å$^2$.

DNA-clamping state:
19,619 micrograph movies were collected on eBIC Titan Krios III coupled to a Falcon VI detector and Selectris X energy filter, with an operating acceleration voltage of 300 keV, a nominal magnification of 130k and pixel size of 0.931 Å, at a dose rate of 12 $e^{-}$/Å/s for a total overall dose of 40 $e^{-}$/Å$^2$.

DNA-clamping state with M18BP1:
18,877 micrograph movies were collected on a Glacios 2 coupled to a Falcon IV detector and Selectris X energy filter, with an operating acceleration voltage of 200 keV, a nominal magnification of 165k and pixel size of 0.69 Å, at a dose rate of 13 $e^{-}$/Å/s for a total overall dose of 50 $e^{-}$/Å$^2$.

Cryo-electron microscopy data processing

Apo condensin II:
Micrograph movies were imported into RELION [71], where motion correction was performed using RELION's own implementation and contrast-transfer function (CTF) parameters estimated, using CTFFIND4 [72]. Particles were picked automatically using cryolo [73], then imported into RELION for extraction. Particles were extracted (4x binning, box size 120 pixels) and exported into cryoSPARC [74]. Iterative 2D classification was performed (100 classes, 18 Å resolution limit) and classes bearing clear domain features were used for *ab initio* model generation (3 classes, 18 Å resolution limit). Putative condensin II volumes were then subject to one round of homogeneous refinement, using default parameters, followed by three rounds of heterogeneous refinement (6 classes, 18 Å resolution limit) with one condensin II dimer volume, one hinge volume, and four junk decoy volumes computed from selected 2D classes with no obvious protein features. Particles classifying into the condensin II dimer volume were exported into RELION using Pyem, and a single round of 3D Refinement was performed using default parameters, and with imposition of C2 symmetry. Following a single round of refinement, iterative CTF Refinement and Bayesian particle polishing were performed, followed by a final round of 3D Refinement. At this stage, processing for the condensin II dimer was considered complete. To achieve more homogeneous volumes of the dimerization and NCAPH2$^{\mathrm{Neck}}$-

NCAPG2 interfaces, the processing diverged into two streams, in which the particles were first subjected to symmetry expansion with RELION and C2 symmetry.

Apo condensin II: Neck stream.
Symmetry-expanded particles were subjected to particle subtraction (RELION) using a mask around the NCAPD3-NCAPG2-putative NCAPH2 density, this was followed by a further round of focused 3D Refinement, and a final round of 3D classification (3 classes, no alignment, T=324) that resulted in the final volume used for model building.

Apo condensin II: Dimerization interface stream.
Symmetry-expanded particles were subjected to particle subtraction (RELION) using a mask around the NCAPD3-SMC4-NCAPH2 and the N- and C-termini of NCAPG2 originating from protomer monomer 1, and 2, respectively. A further round of 3D classification was performed (3 classes, no alignment T=24) followed by 3D Refinement, and a single round of MultiBody refinement with two masks: one contoured around the NCAPD3-SMC4-NCAPH2-NCAPG2$^{\text{Protomer1-N-terminus}}$, and the other around the SMC4-NCAPH2-NCAPG2$^{\text{Protomer2-C-terminus}}$).

ATP-engaged autoinhibited state:
Micrograph movie motion correction, CTF estimation and 2D classification was performed on-the-fly using cryoSPARC Live. Initial blob picking and 2D classification were used to generate templates with clear proteinaceous features for template-based picking, also in cryoSPARC Live. Particles were then exported into cryoSPARC. We initially performed a data processing strategy identical for the apo condensin II complex, however ultimately higher resolution reconstructions were recovered when following a pathway in which selected 2D classes from cryoSPARC Live processing were used directly for *ab initio* model generation, followed by homogeneous refinement. Particles corresponding to the ATP-engaged condensin II were then subjected to one round of homogeneous refinement, using default parameters, followed by non-uniform refinement with CTF correction enabled. Particles were then subjected to reference-based motion correction and exported to RELION. A single round of consensus 3D Refinement was performed, followed by particle subtraction using a mask excluding the flexible termini of NCAPD3-NCAPG2, and the SMC2-SMC4 coils. A final round of 3D Refinement was then performed.

Activated M18BP-bound condensin II:
Data processing for the activated M18BP1-bound condensin II complex followed procedures identical to those described for the ATP-engaged state up to reference-based motion correction. Following reference-based motion correction and a further round of non-uniform refinement it became apparent that the HAWK subcomplexes were better defined than the ATPase domains of SMC2 and SMC4. The processing strategy therefore diverged between the HAWK focus and the ATPase focus. For the HAWKs, the particles were exported to RELION, subjected to a further round of 3D Refinement, after which signal subtraction was performed using a mask around the HAWK density. The HAWK-retaining particles were then input into a final round of 3D Refinement.

For the ATPase focus, we performed 3D variability analysis in cryoSPARC (3 classes, 8 Å resolution filter) followed by 3D variability display (clustering mode, 8 Å resolution filter). Visual inspection of the map indicated the presence of three species, one in which the ATPase domains were not well defined, another in which the ATPases adopted the ATP-engaged conformation, and another in which the ATPases were non-engaged. Particles corresponding to the ATP-engaged and non-engaged conformations were then independently subjected to a final round of non-uniform refinement with CTF correction enabled.

DNA-clamping state:
Micrograph movies were imported into RELION, where motion correction was performed using RELION's own implementation and contrast-transfer function (CTF) parameters estimated, using CTFFIND4. Particles were picked automatically using cryolo, then imported into RELION for extraction. Particles were extracted (4x binning, box size 120 pixels) and exported into cryoSPARC. Iterative 2D classification was performed (100 classes, 18 Å resolution limit) and classes bearing clear domain features were used for *ab initio* model generation (3 classes, 18 Å resolution limit). Putative condensin II - DNA volumes were then subject to one round of homogeneous refinement, using default parameters, followed by three rounds of heterogeneous refinement (6 classes, 18 Å resolution limit) with one condensin II - DNA volume, and four junk decoy volumes computed from selected 2D classes with no obvious protein features. Particles classifying into the condensin II - DNA volume were exported into RELION using Pyem, and a single round of 3D Refinement was performed using default parameters. Following a single round of refinement, iterative CTF Refinement and Bayesian particle polishing were performed, followed by a final round of 3D Refinement.

DNA-clamping state with M18BP1:
Data processing for the DNA-clamping state with M18BP1 was identical to those applied for the ATP-engaged, and activated M18BP1-bound condensin II complexes except that the resolution of the final reconstruction was deemed insufficient for unbinning and export to RELION.

Model building and refinement

For all atomic models, manual model building was performed in Coot [75] followed by real-space refinement with PHENIX [76].

Apo condensin II: Dimer
AlphaFold3 was used to predict binary structures between the kleisin and other non-kleisin subunits. The NCAPD3-NCAPH2, SMC4-NCAPH2, and NCAPG2-NCAPGH2 predictions were fit as rigid bodies into the dimer map using ChimeraX [77].
The coordinates were then exported into Coot for manual correction, with flexible regions excised from the models, and regions with well-defined side-chains manually corrected using Coot.

Apo condensin II: Neck stream.
The NCAPD3-NCAPG2 interface from the dimer coordinates was fit into the map density arising from the NCAPH2$^{Neck}$-focused stream. The NCAPH2$^{Neck}$ density was automatically built using ModelAngelo [28], whereas density from the NCAPD3$^{Tail}$ were manually traced and built in Coot using the unsharpened map to position the peptide backbone, and the sharpened map to verify the sequence register.

Apo condensin II: Dimerization interface stream.
The NCAPD3-NCAPH2-SMC4$^{Protomer1}$ and NCAPG2$^{Protomer2}$ coordinates were fit as rigid bodies in ChimeraX into the dimerization focused map density. Manual correction was then performed in Coot, with poorly defined regions excluded from the models.

ATP-engaged autoinhibited state:
NCAPD3-NCAPH2, SMC2, SMC4-NCAPH2, and NCAPG2-NCAPGH2 AlphaFold3 predictions were fit as rigid bodies into the monomer map using ChimeraX. The coordinates were then exported into Coot for manual correction, with flexible regions excised from the models, and regions with well-defined side-chains manually corrected using Coot. ATP-$Mg^{2+}$ was placed into the map manually.

ATP-engaged M18BP1 activated state:
The ATP-engaged autoinhibited ATPase subcomplex containing SMC2-SMC4-NCAPH2 was rigid body fitted into the map using ChimeraX. The NCAPD3-NCAPH2, and NCAPG2-NCAPH2-M18BP1 AlphaFold3 predictions were separately fit as rigid bodies into the ATP-engaged activated map using ChimeraX. The coordinates were then exported into Coot for manual correction, with flexible regions excised from the models, and regions with well-defined side-chains manually corrected using Coot. ATP-$Mg^{2+}$ was retained in the map based on superpositioning with the higher-resolution ATP-engaged autoinhibited state.

DNA-clamping state:
The ATP-engaged autoinhibited ATPase subcomplex containing SMC2-SMC4-NCAPH2 was rigid body fitted into the map and manually adjusted using Coot. The SMC2$^{Neck}$:NCAPH2$^{Neck}$ was replaced with an AlphaFold3 prediction of the corresponding domains. The NCAPD3-NCAPH2 subcomplex was rigid-body fitted into the map, followed by manual correction. A DNA duplex was similarly manually placed into the density using Coot and adjusted by real-space refinement in Coot.

<u>Protein complex purification for biochemistry and single molecule experiments</u>

Purification of the NCAPG2-NCAPH2 subcomplex:
NCAPG2-NCAPH2(313-415)-TEV-StrepII were co-expressed in *Trichoplusia ni* (Hi5) insect cells (2 L culture). Cells were harvested by centrifugation at 2,000 × g for 15 min at 20 °C, resuspended in 1× PBS, pelleted again, flash-frozen in liquid nitrogen, and stored at −80 °C. Cell pellets were resuspended in five pellet volumes of lysis buffer (300 mM NaCl, 40 mM HEPES pH 7.8, 10% glycerol, 1 mM EDTA) supplemented with protease inhibitor cocktail (Roche) and Benzonase (Invitrogen). Cells were lysed on ice by sonication (70% amplitude, 4 s on/8 s off for 7 min, followed by 75% amplitude for 1 min). Lysates were clarified by centrifugation at 16,000 rpm for 1 h at 4 °C, and the supernatant was filtered through a 0.5 µm filter.

The filtered lysate was loaded onto a Streptactin affinity column (Qiagen) pre-equilibrated with lysis buffer. After washing, bound protein was eluted with lysis buffer supplemented with 5 mM desthiobiotin (IBT). Eluted fractions were diluted to 150 mM NaCl and applied to a Heparin column equilibrated in buffer containing 150 mM NaCl, 40 mM Tris pH 7.8, 10% glycerol, 1 mM EDTA, and 1 mM DTT. Protein was eluted using a step elution with 500 mM NaCl in the same buffer.

Peak fractions were concentrated and further purified by size-exclusion chromatography on a HiLoad 16/600 Superose 6 pg column (Cytiva) equilibrated in SEC buffer (150 mM NaCl, 20 mM HEPES pH 7.8, 5% glycerol, 1 mM TCEP). Fractions were analyzed by SDS–PAGE, pooled, concentrated, flash-frozen in liquid nitrogen, and stored at −80 °C.

Purification of condensin II:
Condensin II complexes for single-molecule experiments were purified similarly to that previously described [19]. Briefly, condensin II subunits were assembled into biGBac vectors[78], transposed into DH10EMBacY cells and purified bacmid transfected into SF9 insect cells. Virus was harvested after 72 hours after transfection and amplified for 72-120 hours, before harvesting and used for expression in SF9 cells for 72 hours. Infected SF9 cells were lysed in condensin purification buffer (20 mM HEPES [pH 8], 300 mM KCl, 5 mM MgCl2, 10% glycerol) supplemented with 5 mM beta-mercaptoethanol, 1 Pierce protease inhibitor EDTA–free tablet (Thermo Scientific) per 50mL and 25 U/mL of Benzonase (Sigma) and cleared lysate loaded onto either a StrepTrap HP or StrepTrap XT (Cytiva) column, washed with condensin buffer before being eluted in condensin buffer supplemented with 10 mM Desthiobiotin or 50 mM biotin, respectively. Condensin complexes were further purified using HiTrap Heparin HP column (cytiva), run with buffer A (20 mM HEPES [pH 8], 150mM NaCl, 5 mM MgCl2, 5% glycerol, 1 mM DTT), eluted with buffer A with 500 mM NaCl and size exclusion chromatography using a Superose 6 increase 10/300 column run with Condensin purification buffer supplemented with 1 mM DTT. Purified condensin II complexes were analyzed by SDS page and mass photometry.
Condensin II pentamers harbouring mutations in NCAPD3 were either expressed from one virus or from two viruses (tetramer lacking NCAPD3 and mutant NCAPD3) with comparable homogeneity, as determined by mass photometry.

M18BP1 residues 874-1132 were cloned into modified pET vector, with N-terminal 10xHis tag and C-terminal ybbR and MBP tag. Protein was expressed using BL21 cells, inducing with 0.5 mM IPTG overnight at 18 degrees. Cells were lysed in M18BP1 buffer (20mM Hepes, pH 7.5, 200mM NaCl, 10% glycerol) supplemented with 10 mM imidazole and 5 mM beta-mercaptoethanol via sonication and cleared lysate applied to TALON superflow resin (Cytiva). Resin was washed with M18BP1 buffer supplemented with 20 mM imidazole, before eluting with M18BP1 buffer 500 mM imidazole. M18BP1 was further purified using HiTrap Heparin HP column (cytiva), run with buffer A (20 mM HEPES [pH 8], 150mM NaCl, 5 mM MgCl2, 5% glycerol, 1 mM DTT) and size exclusion chromatography using condensin purification buffer and a Superose 6 or Superdex 200 increase 10/300 column.

Condensin II and M18BP1 were fluorescently labelled using the SFP transferase to couple a CoA conjugated to Alexa647 or JF646, respectively, via an ybbR tag at the C-terminus of M18BP1 or in the N-terminus of SMC4 in Condensin II. Purification of SFP and labelling was performed as previously described [79], except using a 5–fold excess of dye-CoA over protein. Unconjugated dye and SFP was separated from labelled complex using size exclusion chromatography with a Superose 6 increase 10/300 column and the presence specific conjugation confirmed by SDS page.

ATPase assays

ATPase assays were performed as previously described [17], using the EnzChek Phosphate Assay Kit (Invitrogen) modified for a 96 well plate format [80], with a final protein concentration of 50 nM and a final salt concentration of 100 mM KCl. Absorbance at 360 nm was read using a kinetic protocol on FLUOstar omega (BMG Labtech) plate reader and phosphate release rate calculated using a standard curve of known phosphate concentration.

Electromobility Shift Assays

Binding reactions contained NCAPG2-NCAPH2 alone, M18BP1 wild-type (wt) alone, M18BP1 mutant alone, or NCAPG2-NCAPH2 pre-mixed with M18BP1 (wt or mutant). For mixed samples, NCAPG2-NCAPH2 and M18BP1 were combined at a 1:1.2 molar ratio prior to DNA addition. Proteins (0.5–4 µM final concentration) were incubated with 1 µM of a 180-bp DNA fragment in binding buffer (100 mM NaCl, 20 mM HEPES pH 7.8, 5% glycerol, 1 mM TCEP) in a total volume of 10 µL for 30 min at room temperature. Reactions were supplemented with 2 µL 6× NEB Gel Loading Dye (Purple) and resolved on a 1% agarose gel in TBE buffer at 88 V for 25 min. Gels were stained with SYBR Safe to visualize free DNA and DNA–protein complexes.

Cell lines and constructs

Chicken DT40 CDK1$^{as}$ cells were cultured in RPMI 1640 medium (Thermo Fisher Scientific, catalog no. 21875091) supplemented with 10% FBS and 1% chicken serum, and maintained at 39 °C in a humidified incubator with 5% $CO_2$. OsTIR1$^{F74G}$ cDNA under the control of the CMV promoter with G418 resistance cassette was, randomly integrated into the genome to generate DT40 CDK1$^{as}$/OsTIR1$^{F74G}$ cells, which constitutively express OsTIR1$^{F74G}$, as previously described[1].

To establish CAPD3-AID cells, a rescue construct (~500 bp homology arms for the C-terminus of GgCAPD3 gene and a miniAID-Clover tag with a hygromycin resistance cassette), and a PX330-derivative plasmid (a guideRNA targeting *GgCAPD3* and Cas9) was transfected into the DT40 CDK1$^{as}$/OsTIR1$^{F74G}$ cells. PX330 is a gift from F. Zhang (Addgene plasmid 42230). Double stranded oligos used to target *GgCAPD3* are aggtcccagaccgtgctc, gccatccgtgccacggga. Establishment of CAPD3-AID cells were verified using an anti-GgCAPD3 antibody (a gift from D. Hudson[81]). Depletion of the CAPD3 protein was confirmed by flowcytometry analysis.

Doxycycline-inducible expression of wild-type or mutants *GgCAPD3* fused at the C-terminus to mScarlet3 and V5 tag was achieved by transient transfection of plasmids derived from pMK406 (a gift from M. Kanemaki, NIG, JAPAN) and a plasmid encoding the PiggyBac transposase. PX330-derivative plasmids targeting *GgMCPH1* or *GgMis18BP1* were constructed using oligos Gg*MCPH1* (caccgggagaaaagcacaggatgc, aaacgcatcctgtgcttttctccc) and Gg*Mis18BP1* (caccggatcagcgagatccgcccc, aaacggggcggatctcgctgatcc), respectively.
Plasmids were constructed by K Samejima using restriction digest and the maps for rescue construct and for CAPD3 expression are provided separately.

Premature Chromosome Condensation Assays

45-47h prior to harvesting cells, ~3X10$^{6}$ CAPD3-AID cells were transfected with plasmids coding wild type or mutant CAPD3-mScarlet-V5 (~2 µg), PiggyBac transposase (~1 µg), and

PX330 derivative plasmids (~5 µg of either control, targeting *GgMIs18BP1*, and/or *GgMCPH1* to inactivate corresponding genes) using 100 µl Neon NxT Electroporation system (buffer R, voltage 1600, Pulse width 10, Pulse number 3) and resuspended into 10 ml culture media. At –13h, doxycycline (final concentration 0.5 µg/ml) and 1NMPP1 (final concentration 2 µM) were added to the culture to induce the expression of wild-type or mutants GgCAPD3. At –3h, 5Ph-IAA (final concentration 10 µM) were added to the media to acutely deplete endogenous CAPD3 protein and then 1 ml of cells were plated on poly-L-lysine coated coverslips (Fisher scientific 10468681) in 16 well dishes.

At 0h, cells plated on coverslips were rinsed with prewarmed PBS and fixed with 4% Formaldehyde (Thermofisher Scientific catalog No. 28908) in PBS for 10 min. Cells were stained with DNA dye (Hochest 33452) and mounted with Vectashield plus antifade mounting medium (Vector-lab H-1900). Images of >50 scarlet positive cells were taken with a wide-field Deltavision Elite microscopy system [Olympus iX71 Inverted Fluorescence Microscope, SoftWoRx software 7.0.0(Applied Precision Inc, Image Solutions UK Ltd), PCO edge 4.2 sCMOS camera, a X 100 UPLX-Apochromat objective/1.45 Oil]. 3D datasets (11 Z sections) were visualized and analyzed using Fiji. Number of cells exhibiting Premature Chromosome Condensation (PCC), those showing some degree of condensed chromosomes (PCC-like), or those with uncondensed interphase chromatin (normal) were manually counted. Number of biological replicates for $NCAPD3^{WT}$, $NCAPD3^{\Delta Tail}$ and $NCAPD3^{\Delta Tail\text{-}H2}$ are 5, 4, and 4 respectively.

Mass photometry experiments

The mass photometry experiments were performed on a TwoMP device (Refeyn) using the coverslips and silicone gaskets from sample prep kit MP-CON-21022 (Refeyn). The data acquisition was done for 60 s at 128.2 Hz for a large field of view. All of the buffers used in mass photometry experiments were filtered with a 0.22 µm syringe filter (polyvinylidene difluoride membrane, Merck Millex). Measurements were done with a buffer containing 40 mM Tris-HCl pH 7.4, 100 mM Potassium Glutamate, 5 mM $MgCl_2$ and 1 mM TCEP (Figure 1F and S5F) or 50 mM Tris pH 7.4, 150 mM KCl, 5 mM $MgCl_2$ (Figure S4G). The mass photometry buffers for the measurements with ATP were supplemented with the indicated concentrations of ATP (a range of 1 µM to 2.5 mM). The measurements were performed for 5-10 nM of final condensin II concentration. For the mass photometry experiments comparing condensin II-binding of M18BP1, $M18BP1^{D3\ mut}$ and $M18BP1^{NC}$ a final concentration of 5 nM of condensin II and 20 nM of the respective M18BP1 was used. The data was acquired with the AcquireMP (Refeyn) and analyzed with DiscoverMP (Refeyn) softwares. For the contrast to mass conversion Thyroglobulin or MassFerence P1 Calibrant (MP-CON-41033, Refeyn) were used to create a calibration curve on the day of the measurement.

Single molecule assay

The single molecule experiments were carried out as previously described[82], with some minor modifications.

Preparation of biotinylated DNA substrates:

The 44.24 kbp DNA substrates biotinylated at both ends were prepared using linearized cosmid-I95[30]. The cosmid was amplified in NEB 10-beta, purified using QIAfilter Plasmid Midi Kit (QIAGEN) and linearized supping SpeI-HF (NEB) restriction enzyme for 3 hours at 37°C. The biotinylated handles were generated with a PCR reaction on pBluescript SK+ with GoTaq Polymerase (Promega) with the primers 306 bp for (ACGCCAGGGTTTTCCCAG) and 306 rev (TCCGGCTCGTATGTTGTGTG) in the presence of 1/5 Bio-dUTP (Jena Bioscience) to Bio-dTTP and digested using SpeI-HF (NEB) restriction enzyme for 1 hour at 37°C. The handles were purified with Wizard® SV Gel and PCR Clean-Up System (Promega) resulting in 154 bp and 150 bp biotinylated handles. The linearized cosmid and handles were ligated with T4 DNA Ligase (NEB) overnight at 16°C. The resulting 44.24 kbp DNA with biotinylated handles was separated from the handles with a 1% agarose TAE gel and eluted with Roti ® elution Tubes MIDI (Roth).

Functionalization of glass slides and flow cell assembly:

The microscope slides (76x26x1 mm, Marienfeld 1000000) were laser drilled for the attachment of inlet/outlet tubings and reused multiple times. The microscope slides were cleaned by sonication with Citranox detergent (15 minutes), MQ water (10 minutes), acetone (15 minutes) and MQ water (5 minutes) after each use. The microscope slides and coverslips (24x60 mm No1.5 thickness 170 µm, borosilicate VWR Intl 631-0147) were then cleaned by sonication in 1 M KOH solution for 30 minutes and treated with acid piranha solution (5:1 sulfuric acid: hydrogen peroxide) for an hour. After piranha etching the microscope slides and coverslips were washed and sonicated (10 minutes) with methanol. Afterwards the slides were silanized for 30 minutes in a 10% APTES and 5% acetic acid solution in methanol. After the silanization the slides were washed extensively with methanol and dried under a flow of compressed nitrogen. Finally, the slides were PEGylated with 40 mg/ml of m-PEG-SVA (5000 Da Laysan Bio) and 1 mg/ml of Biotin-PEG-SVA (5000 Da Laysan Bio) in 50 mM Boric Acid pH 8.5. The PEGylation step was repeated for 4 times for at least 4 hours at 4°C. After the 4$^{th}$ PEGylation step the slides were PEGylated with shorter PEG molecules, 50 mM MS(PEG)4 (Thermo Scientific Pierce) in 50 mM Boric Acid pH 8.5. In between each washing step the slides were washed thoroughly with MQ water and dried under a flow of compressed nitrogen. After the last PEGylation step the slides were dried and sealed with nitrogen and stored at -20°C until use.

A flow cell was assembled with one microscope slide and coverslip, using thinly cut double sided scotch tape as a spacer between the two to generate 2 mm x 24 mm x 100 µm channels. The channels were then sealed with Epoxy glue. Tubings were attached to the flow cell by 200 µl pipette tips and used as outlets. The flow was controlled by a syringe pump, attached to the tubing by a needle on its other end. 200 µl pipette tips were used as inlets during experiments.

HiLo microscopy and data acquisition:

The imaging of the single molecule experiments was done on a custom built highly inclined optical light sheet (HiLo) microscope, as described previously in Pradhan et al[82]. Briefly, 561 nm and 638 nm wavelength lasers were coupled to an optical fiber and directed into an AxioVert 200 Zeiss microscope with an adjustable mirror to change the angle of illumination. A total internal reflection fluorescence (TIRF) objective (Objective alpha Plan-Apochromat 63x/1.46 Oil) was used. The illumination was adjusted for HiLo mode, where DNA and proteins directly above the surface were selectively illuminated by an optical light sheet, while supressing the background intensity from the fluorophores inside the solution. For the simultaneous imaging of Sytox Orange labelled DNA and Alexa Fluor 647 or Janelia Fluor 646 labeled proteins an alternative excitation (ALEX) was used. A custom developed software in napari and pyqtgraph was used for the operation of the sCMOS camera (PCO edge 4.2) and the lasers, for image acquisition and visualization.

Single molecule loop extrusion assay:
The flow cell assembled as described above was used for the single molecule assay for the visualization of loops formed by condensin II. Each channel was first washed thoroughly with T50 buffer (40 mM Tris-HCl pH 8.0, 50 mM NaCl 0.2 mM EDTA), then incubated with 1 µM Neutravidin in T50 buffer for 2 minutes and 5 mg/ml ultrapure BSA in T50 buffer for 5 minutes. The excess Neutravidin and BSA were washed away with T50 buffer. 10-30 pM of linear 44.24 kbp DNA biotinylated at both ends was introduced to the channel with a flow rate of 2 µl/min and tethered to the surface by the biotin-Netravidin binding. The flow rate was adjusted and kept constant by a syringe pump to achieve a relative extension of 0.3-0.4. Excess DNA not bound to the surface was washed away with T50 buffer.

The DNA tethered on the surface was visualized with the imaging buffer containing 40 mM Tris-HCl pH 7.4 37°C, 2 mM Trolox, 100 mM Potassium Glutamate, 5 mM $MgCl_2$, 30 mM D-Glucose, 1 mg/ml BSA, 1 mM TCEP, 2.5 mM ATP, 100 nM Sytox Orange, 0.15 mg/ml Glucose Oxidase and 0.02 mg/ml Catalase. The imaging buffer not containing the protein was flushed into the channel to allow for equilibration directly prior to imaging and the flow cell was heated to 37°C and kept at 37°C for the image acquisition. The imaging buffer supplied with protein, the concentration details as described below, was introduced to the channel with a flow rate of 10 µl/min for 100s and the flow was stopped afterwards. Typically, a measurement lasted 1000s and 10000 frames were acquired.

In the experiments with labeled condensin II, 3-10 nM of Alexa Fluor 647 labelled condensin II was used. Loop-extrusion activity was quantified as looping probability, defined by the fraction of DNA molecules that formed at least one loop during a 1000-second acquisition period. For the comparison of looping probability of condensin II mutants, 2 nM of each condensin II variant (WT, ΔG2, $D3^{\Delta Tail}$, $D3^{\Delta Tail\text{-}G2}$ and $D3^{Tail\text{-}H2\ Ala}$) was used. For the comparison of looping probability of condensin II in the presence of M18BP1, 2 nM condensin II and various concentrations of M18BP1 (2 nM, 8 nM, 16 nM, 50 nM, 100 nM and 200 nM) was used.
For the experiments where condensin II and M18BP1 were phosphorylated in vitro, a 25x molar excess of M18BP1 to condensin II was used for non-labeled M18BP1 and 5x molar excess of M18BP1 to condensin II was used for Janelia Fluor 646 labeled M18BP1. Condensin II and M18BP1 were incubated at room temperature for 30 minutes together with CDK1/Cyclin B in the presence of 2 mM ATP directly prior to imaging.

For the comparison of looping probability of pre-phosphorylated samples ($WT^{phos}$, $M18BP1^{phos}$, $WT^{phos}$+$M18BP1^{phos}$) 2 nM of each protein was used. For the comparison with these samples, another batch of condensin II WT, purified under the same conditions was used, yielding a similar looping probability as the other purification conditions.

For the experiments with Janelia Fluor 647 labelled M18BP1, 10 nM of M18BP1 was used either alone, in the presence of 2 nM condensin II WT, 2 nM condensin II WT and 5 nM CDK1 and 10 nM Cylin B or 2 nM condensin II ΔG2.

For the measurements where the loop durations were extracted, the condensin II and M18BP1 concentrations were titrated down, while keeping the molar ratios constant, to avoid multiple loop events on the same DNA strand.

Data Analysis:

Analysis of acquired single molecule image series was done with a custom written software in Python, based on the packages Napari (https://github.com/Napari/napari) and PyQtGraph (https://github.com/pyqtgraph/pyqtgraph). The individual details are described below. For a single dataset, a field of view of approximately 92 µm x 92 µm containing around 100-200 DNA strand was acquired and analysed. The datasets were inspected visually for identification of loops. A loop was defined as a puncta of higher fluorescence intensity on a DNA strand with gradually increasing intensity. The snapshots shown in figures were smoothed with a median filter of 2 pixel radius and background subtracted with a white top-hat filter with 10 pixel radius. The kymographs were built by the sum of fluorescence intensity of 15 pixels along the lines perpendicular to the line connecting the two DNA tethering points. For the datasets where DNA and labelled condensin II were imaged simultaneously, separate kymographs were built from the corresponding image sequences recorded using ALEX.

*Analysis of bleaching steps, number of fluorophores and colocalization of loop start with protein signal for labelled condensin II and labelled M18BP1*

The DNA and protein kymographs of loops formed in the presence of Alexa Fluor 647 were inspected manually for colocalization of protein signal with loop start. The intensity time traces were calculated using the sum of intensity of the protein signal in a line of 9 pixels around the intensity maxima for each time point. The bleaching steps were inferred from the intensity time traces. The loops where the protein signal could not be clearly assigned to a number of bleaching steps, due to e.g. overlapping signal with surface stuck molecules, or low protein signal due to the non-even illumination, were discarded.

For the estimation of the number of fluorophores of non-looping, DNA-bound condensin II complexes, random DNA strands without any loop were chosen. The protein kymographs were inspected manually for fluorophore signal bound to DNA. The number of fluorophores from a protein signal was estimated by comparison of signal intensity with the signal of proteins stuck to surface non-specifically that bleached in 1 or 2 steps.

DNA and protein kymographs were manually inspected for the estimation of fraction of loops that colocalized with protein signal that were formed in the presence of M18BP1. For the estimation of non-looping, DNA bound M18BP1 signal random DNA molecules without a loop were chosen and manually inspected for number of binding events by a protein signal.

*Analysis of protein looping probability*

To assess the activation of condensin II, “looping probability” was used as a measure of looping activity. Looping probability was defined as the fraction of DNA strands that has at least one loop in the 1000s acquisition time. The fraction of looped DNA was calculated from the large field of view counting the DNA strands with or without loops.

*Analysis of loop kinetics, calculation of loop extrusion rates and loop durations*

For the analysis of loop kinetics the size of DNA regions within and outside the loop were calculated from the fluorescence intensity. For this, each line in a kymograph was split into three regions: “Loop”, “Up” and “Down”, denoting the loop region and the DNA portions above and below it. The center position of the loop was determined from the intensity maxima for each time point and the fluorescence intensity was calculated by summing the intensity of 9 pixels around the loop center. The size of DNA within each region was calculated by the formula:

$$DNA\ size\ (region) = \frac{Intensity\ (region) \times 44240\ kbp}{Total\ DNA\ intensity}$$

The loop extrusion rate at the start phase of a loop was calculated by fitting a linear function $f(x) = kx + c$ to the initial 5 seconds of the loop size.

The loop durations were calculated for the loops from the start of puncta forming to either the end of the loop signal or the end of data acquisition. Two loops formed simultaneously on the same DNA strand were not considered, as the kinetics of one could affect the other.

*Quantification and statistical analysis*

The single molecule data was analysed and visualized with Python, using mainly the packages Numpy (https://numpy.org), SciPy (https://scipy.org) and Matplotlib (https://matplotlib.org). The traces in Fig 1 H were smoothed using a Saviztky-Golay filter in Scipy using 15 datapoints. The traces in Fig 4D,F and Fig S9B,C were smoothed with a Saviztky-Golay filter with 30 datapoints for the WT traces and 100 datapoints for the WT+M18BP1+CDK1 traces.

### Supplemental Information

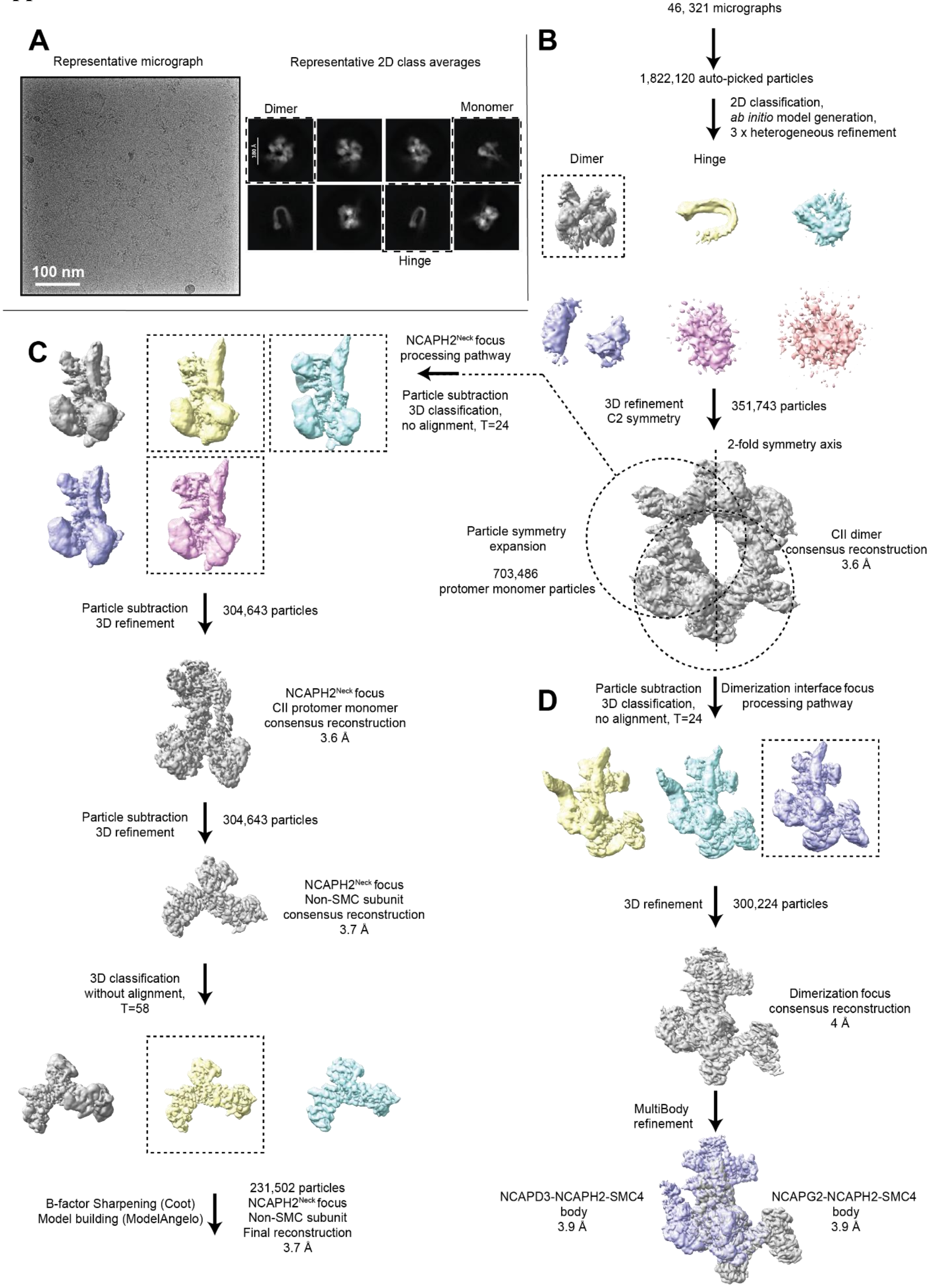

**Figure S1. Workflow for the apo condensin II complex cryo-EM reconstructions. (A)** Representative micrograph and 2D class averages for apo condensin II. **(B)** Workflow for the apo condensin II dataset leading to consensus reconstruction of the condensin II dimer **(C)** Workflow for the NCAPH2$^{Neck}$ focused processing. **(D)** Workflow for the condensin II dimerization interface focused processing.

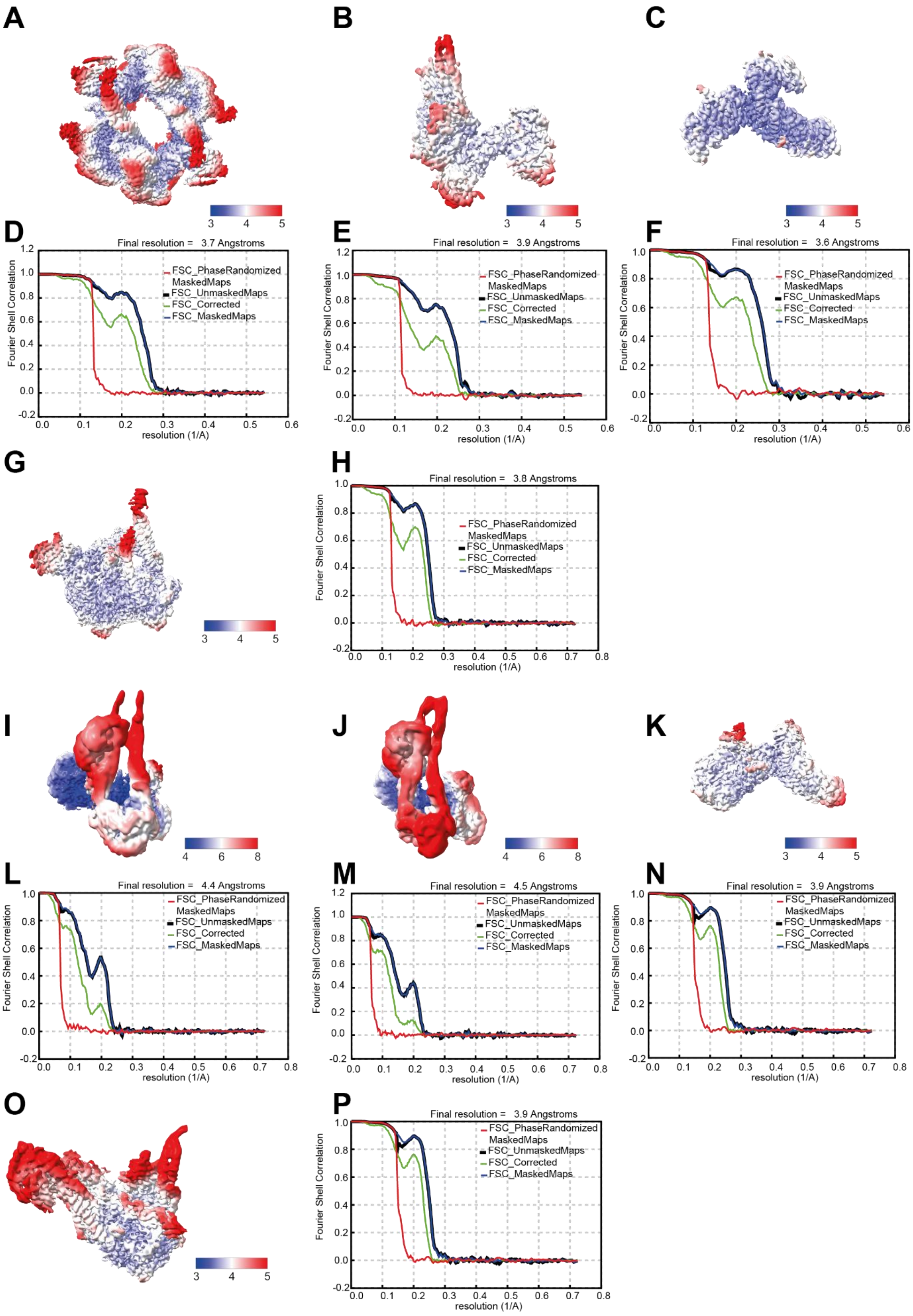
A
B
C
D
Final resolution = 3.7 Angstroms
E
Final resolution = 3.9 Angstroms
F
Final resolution = 3.6 Angstroms
G
H
Final resolution = 3.8 Angstroms
I
J
K
L
Final resolution = 4.4 Angstroms
M
Final resolution = 4.5 Angstroms
N
Final resolution = 3.9 Angstroms
O
P
Final resolution = 3.9 Angstroms
FSC_PhaseRandomized MaskedMaps
FSC_UnmaskedMaps
FSC_Corrected
FSC_MaskedMaps
Fourier Shell Correlation
resolution (1/A)

**Figure S2. Local resolution estimation and FSC curves for cryo-EM reconstructions.** Local resolution estimation of the condensin II dimer **(A)**, dimerization interface focus protomer monomer **(B)**, and NCAPH2$^{Neck}$ focus protomer monomer, and corresponding fourier shell correlation (FSC) curves displayed below **(D-E)**. Local resolution estimation of the ATP-engaged condensin II complex **(G)** and corresponding FSC curves **(H)**. Local resolution estimation of the condensin II – M18BP1 complexes, including the ATP-engaged **(I)**, non-engaged **(J)**, and HEAT-focused **(K)** reconstructions, and corresponding FSC curves displayed below **(L-N)**. Local resolution estimation of the condensin II – DNA – ADP.BeFx complex **(O)** and corresponding FSC curves **(P)**. Map resolution range used is displayed in a color-coded key below the local-resolution filtered maps.

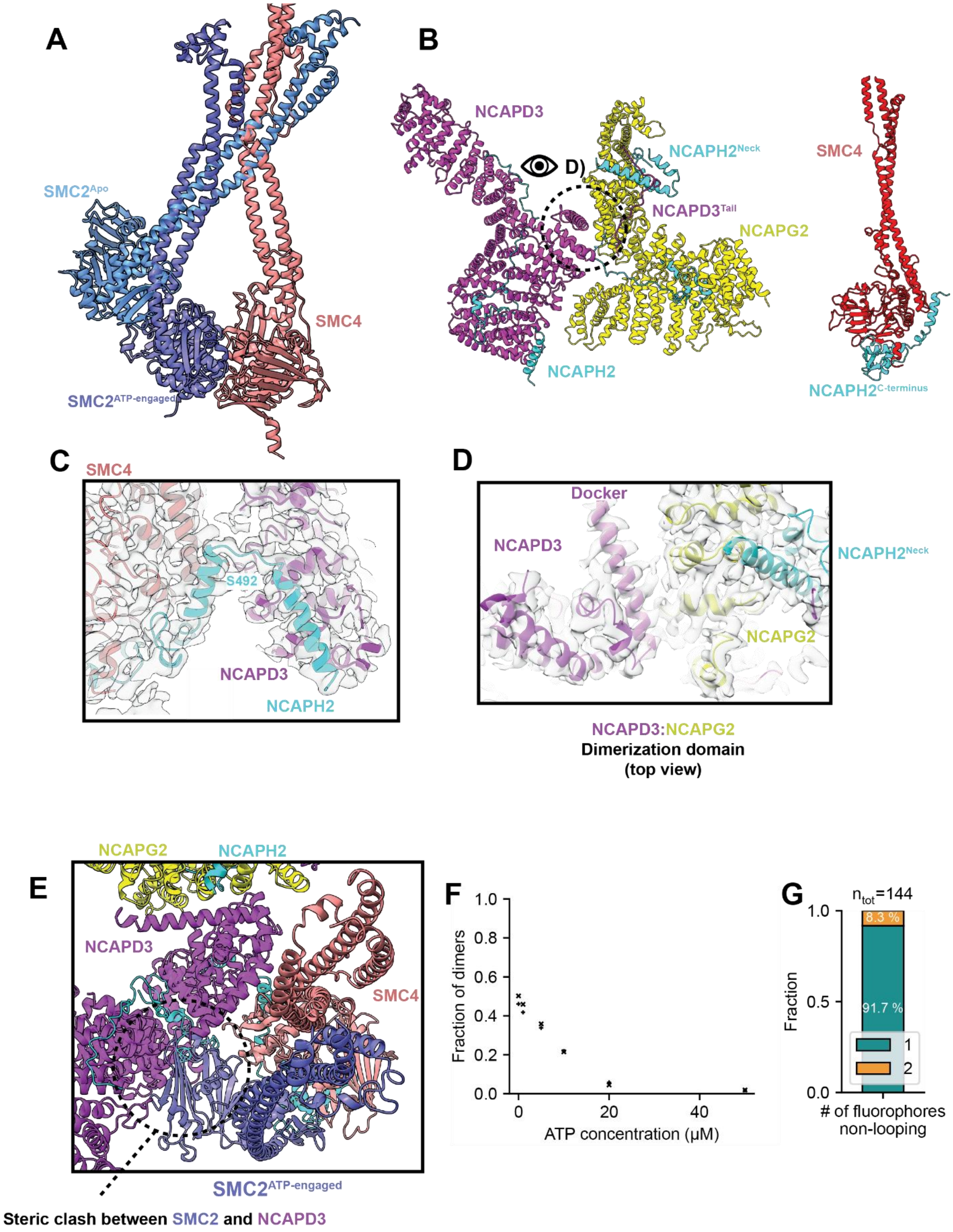

A
SMC2Apo
SMC4
SMC2ATP-engaged
B
NCAPD3
D)
NCAPH2Neck
NCAPD3Tail
NCAPG2
NCAPH2
SMC4
NCAPH2C-terminus
C
SMC4
S492
NCAPD3
NCAPH2
D
Docker
NCAPD3
NCAPH2Neck
NCAPG2
NCAPD3:NCAPG2
Dimerization domain
(top view)
E
NCAPG2
NCAPH2
NCAPD3
SMC4
SMC2ATP-engaged
Steric clash between SMC2 and NCAPD3
F
Fraction of dimers
1.0
0.8
0.6
0.4
0.2
0.0
0
20
40
ATP concentration (μM)
G
ntot=144
1.0
0.5
0.0
Fraction
8.3 %
91.7 %
1
2
### of fluorophores
non-looping

**Figure S3. Additional details of apo condensin II structure and its incompatibility with ATPase engagement.** **(A)** Superpositioning of the apo and ATP-engaged SMC ATPases. **(B)** Structural model of the HEAT-NCAPH2 (left) and SMC4-NCAPH2 subcomplexes (right). **(C)** Cryo-EM density displayed over the SMC4 and NCAPD3 interface bridged by NCAPH2. The position of the S492 Mps1 phospho-site is indicated. **(D)** Cryo-EM density contoured around the NCAPD3-NCAPG2 heterodimerization interface **(E)** Structural model highlighting steric clash introduced between SMC2 and NCAPD3 by ATPase engagement. **(F)** Calculated fraction of condensin II dimers from mass photometry experiments dependent on ATP concentration in a range of 0-50 μM of ATP. Results from two replicates for each condition is included. **(G)** Number of fluorophores per single binding event of DNA-bound non-loop-extruding condensin II molecules labeled with Alexa Fluor 647 dye. $n_{tot}$ = 144.

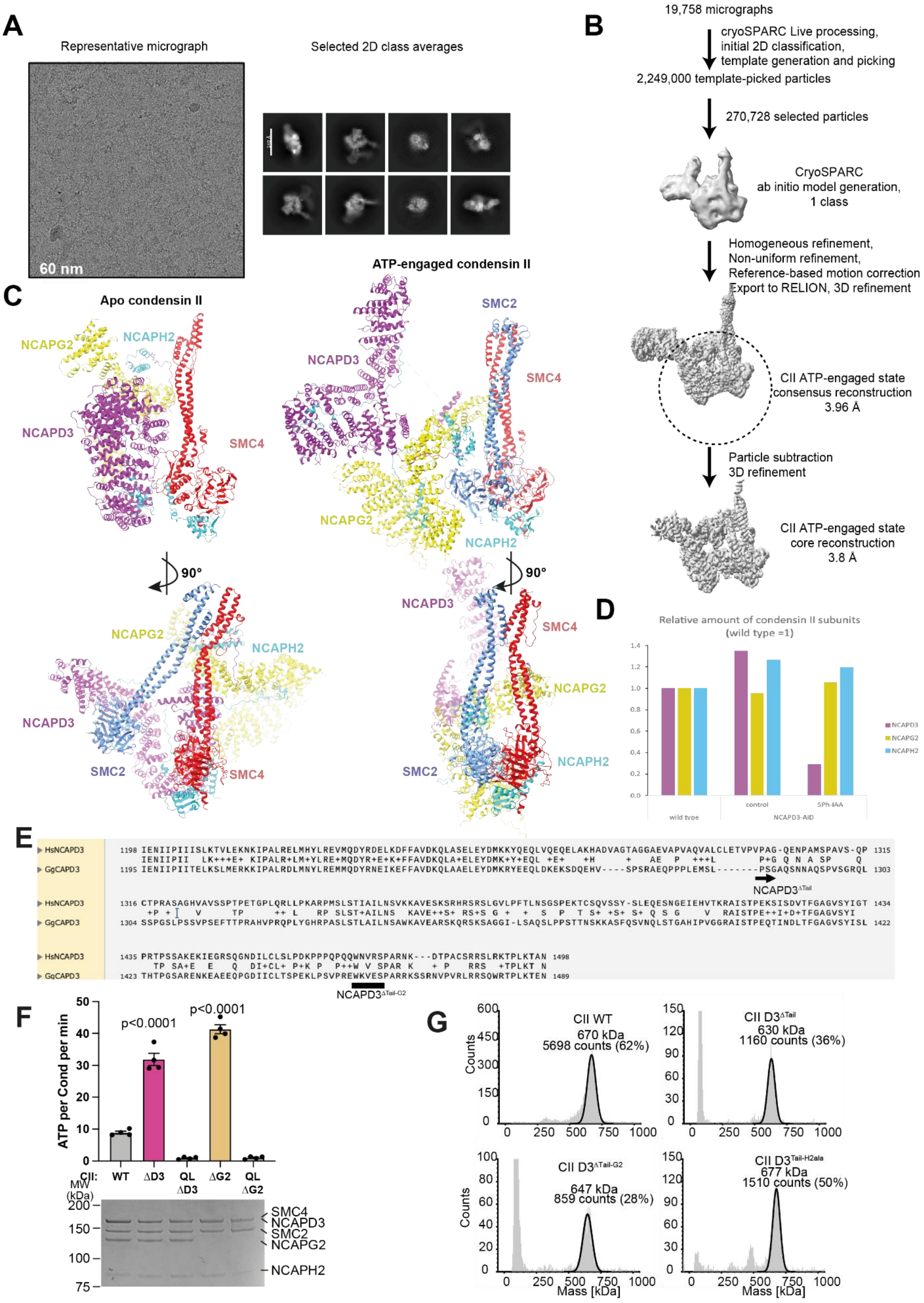
A
Representative micrograph
60 nm
Selected 2D class averages
B
19,758 micrographs
cryoSPARC Live processing,
initial 2D classification,
template generation and picking
2,249,000 template-picked particles
270,728 selected particles
CryoSPARC
ab initio model generation,
1 class
Homogeneous refinement,
Non-uniform refinement,
Reference-based motion correction
Export to RELION, 3D refinement
CII ATP-engaged state
consensus reconstruction
3.96 Å
Particle subtraction
3D refinement
CII ATP-engaged state
core reconstruction
3.8 Å
C
Apo condensin II
ATP-engaged condensin II
NCAPG2
NCAPH2
NCAPD3
SMC4
SMC2
90°
D
Relative amount of condensin II subunits
(wild type =1)
wild type
control
5Ph-IAA
NCAPD3-AID
NCAPD3
NCAPG2
NCAPH2
E
HsNCAPD3
GgCAPD3
NCAPD3ΔTail
NCAPD3ΔTail-G2
F
ATP per Cond per min
p<0.0001
p<0.0001
CII:
WT
ΔD3
QL ΔD3
ΔG2
QL ΔG2
MW (kDa)
SMC4
NCAPD3
SMC2
NCAPG2
NCAPH2
G
CII WT
670 kDa
5698 counts (62%)
CII D3ΔTail
630 kDa
1160 counts (36%)
CII D3ΔTail-G2
647 kDa
859 counts (28%)
CII D3Tail-H2ala
677 kDa
1510 counts (50%)
Counts
Mass [kDa]

**Figure S4. Workflow for the ATP-engaged condensin II complex, mass spectrometric quantification of protein depletion in DT40 cells, biochemical and biophysical analyses of condensin II variants. (A)** Representative micrograph and 2D class averages for ATP-engaged condensin II. **(B)** Workflow for the ATP-engaged condensin II data processing. **(C)** Structural comparison of reorganisation of condensin II between the apo and ATP-engaged states. Superpositioning was guided by SMC4. **(D)** The relative abundance of condensin II non-SMC subunits was quantified by whole-cell mass spectrometry and normalized to the corresponding protein levels in asynchronously growing wild-type (WT) cells. Where indicated, cells were treated with 10 µM 5-Ph-IAA for 3 h prior to harvest to induce NCAPD3 degradation. **(E)** Sequence alignment of Human and Chicken NCAPD3 sequences, indicating the location of the $NCAPD3^{\Delta Tail}$ and $NCAPD3^{\Delta Tail\text{-}G2}$ mutations. **(F)** Quantification of rate of ATP hydrolysis by Wild-type (WT) condensin II, and variants lacking NCAPD3 (ΔD3) or NCAPG2 (ΔG2) subunits. QL signifies variants with SMC2 Q147L, SMC4 Q229L substitutions in the active site unable to bind and hydrolyse ATP. Error bars indicate mean and one standard error. Data from N=4 technical repeats. The P values were calculated using t-tests with Welch's correction. **(G)** Mass photometry of condensin II mutant samples used in Fig. 2F confirming holocomplex integrity.

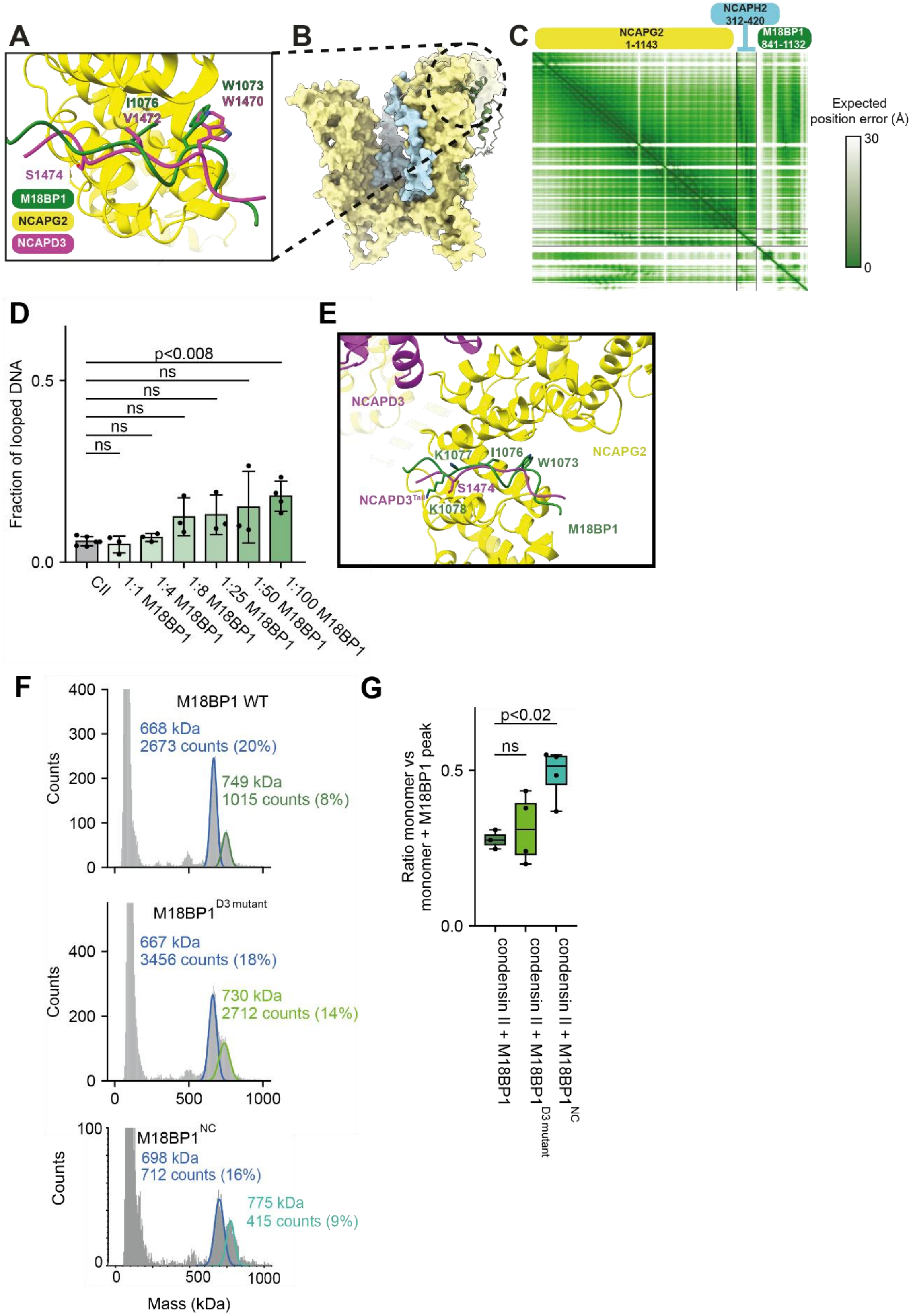
A
W1073
W1470
I1076
V1472
S1474
M18BP1
NCAPG2
NCAPD3
B
C
NCAPH2
312-420
NCAPG2
1-1143
M18BP1
841-1132
Expected
position error (Å)
30
0
D
Fraction of looped DNA
0.5
0.0
p<0.008
ns
ns
ns
ns
ns
CII
1:1 M18BP1
1:4 M18BP1
1:8 M18BP1
1:25 M18BP1
1:50 M18BP1
1:100 M18BP1
E
NCAPD3
K1077
I1076
W1073
NCAPG2
S1474
NCAPD3Tail
K1078
M18BP1
F
M18BP1 WT
668 kDa
2673 counts (20%)
749 kDa
1015 counts (8%)
M18BP1D3 mutant
667 kDa
3456 counts (18%)
730 kDa
2712 counts (14%)
M18BP1NC
698 kDa
712 counts (16%)
775 kDa
415 counts (9%)
Counts
Mass (kDa)
G
Ratio monomer vs
monomer + M18BP1 peak
p<0.02
ns
condensin II + M18BP1
condensin II + M18BP1D3 mutant
condensin II + M18BP1NC

**Figure S5. M18BP1 recognizes a binding site on NCAPG2 that overlaps with NCAPD3$^{Tail}$, and moderately increases looping probability of condensin II (A,B)** AlphaFold3 [42] model showing overlap predicted between NCAPD3 and M18BP1 bound to NCAPG2. **(B)** Prediction confidence of M18BP1 shown green to white, with high confidence (>75%) in green to low confidence in white. **(C)** Expected alignment error of prediction in (B). **(D)** Fractions of DNA molecules exhibiting condensin II-mediated loops in the presence of 2 nM condensin II and 2.5 mM ATP with varying in varying concentrations (1× to 100× excess) of M18BP1. Mean ± s.d. is shown. $n_{tot}$ = 704, 285, 299, 359, 263, 322 and 523. Data are from at least three independent repetitions. The data for condensin II and condensin II 1:100 M18BP1 are the same as in Fig. 3A. The P values were calculated using Welch's t test. **(E)** Structural model of the overlapping NCAPD3$^{Tail}$ and M18BP1 binding site on NCAPG2 with positions of key residues mutated in M18BP1$^{D3}$ annotated **(F)** Exemplary mass photometry histograms of condensin II mass distributions in the presence of M18BP1, M18BP1$^{D3\ mut}$ and M18BP1$^{NC}$ **(G)** Box-whisker-plots showing the fraction of M18BP1-, M18BP1$^{D3\ mut}$- and M18BP1$^{NC-}$bound condensin II monomer, calculated from mass photometry experiments in **(F)**. The central line denotes the median, the box limit denotes the 25th–75th percentile and the whiskers denote minimum and maxima. The P values were calculated using Welch's t test.

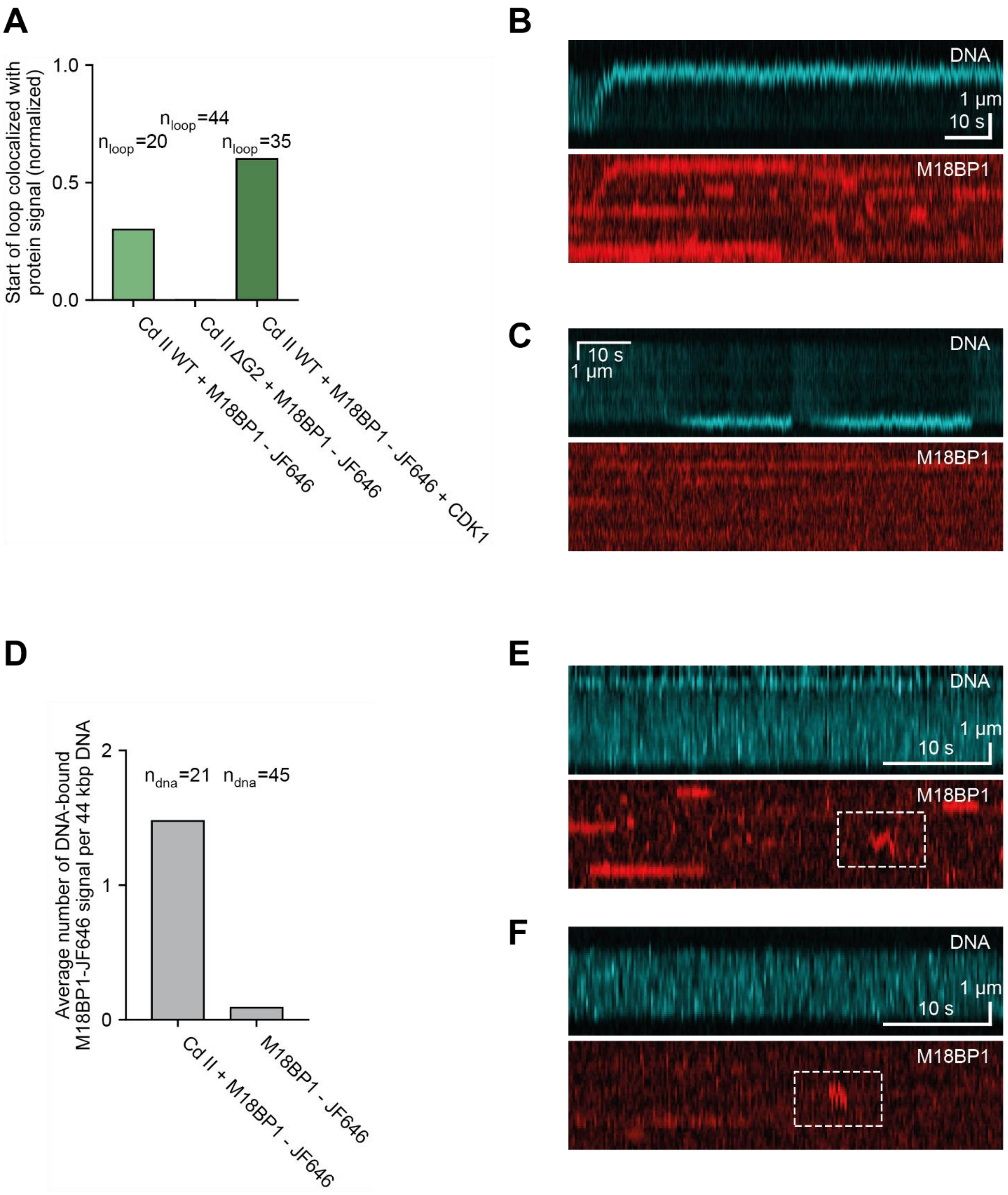
A
Start of loop colocalized with protein signal (normalized)
1.0
0.5
0.0
n_loop=20
n_loop=44
n_loop=35
Cd II WT + M18BP1 - JF646
Cd II ΔG2 + M18BP1 - JF646
Cd II WT + M18BP1 - JF646 + CDK1
B
DNA
1 µm
10 s
M18BP1
C
10 s
1 µm
DNA
M18BP1
D
Average number of DNA-bound M18BP1-JF646 signal per 44 kbp DNA
2
1
0
n_dna=21
n_dna=45
Cd II + M18BP1 - JF646
M18BP1 - JF646
E
DNA
1 µm
10 s
M18BP1
F
DNA
1 µm
10 s
M18BP1

**Figure S6. M18BP1 forms a complex with condensin II capable of loop extrusion: (A)** Fraction of condensin II WT or ΔG2 mediated loops, colocalized with M18BP1 fluorescence signal in the presence 10 nM fluorescence labelled M18BP1 and either 2 nM condensin II WT, 2 nM condensin II ΔG2 or 2 nM condensin II WT and 5 nM CDK1 and 10 nM Cyclin B. The data in the presence of condensin II WT are reproduced from Figure 3E. **(B)** Example kymograph of DNA (top) and M18BP1 (bottom) fluorescence signal of condensin II WT mediated loop showing colocalization of loop start with M18BP1 signal. **(C)** Example kymograph of DNA (top) and M18BP1 (bottom) fluorescence signal of condensin II ΔG2 mediated loop without colocalization of loop start with M18BP1 signal. **(D)** Average number of non-looping, DNA-bound Janelia Fluor 646 labeled M18BP1 fluorescence signal per 44.2 kbp DNA in the presence of 2 nM condensin II and 10 nM M18BP1 or 10 nM M18BP1 in 1000s measurement duration. **(E)** Example kymograph of DNA (top) and non-looping, DNA-bound M18BP1 (bottom) fluorescence signal in the presence of condensin II. **(F)** Example kymograph of DNA (top) and non-looping, DNA-bound M18BP1 (bottom) fluorescence signal in the absence of condensin II.

1035

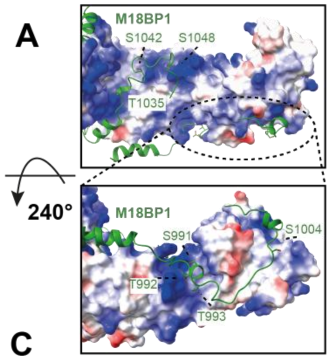

1035

B

| Location | NCAPD3 residue | CDK1 consensus | CII + M18BP1 | CII + M18BP1 + CDK1 | CII + CDK1 |
|---|---|---|---|---|---|
| NCAPG2 interaction | S1348 | no | 1 | 29.4 | 88.8 |
| | S1350 | no | 1 | 3.0 | 6.6 |
| | T1351 | no | 1 | 2.5 | 0.4 |
| | S1357 | no | 1 | 65.7 | 36.3 |
| NCAPH2 interaction | S1458 | almost | 1 | 6.1 | 1.8 |
| M18BP1 competing | S1474 | yes | 1 | 116.1 | 5.6 |

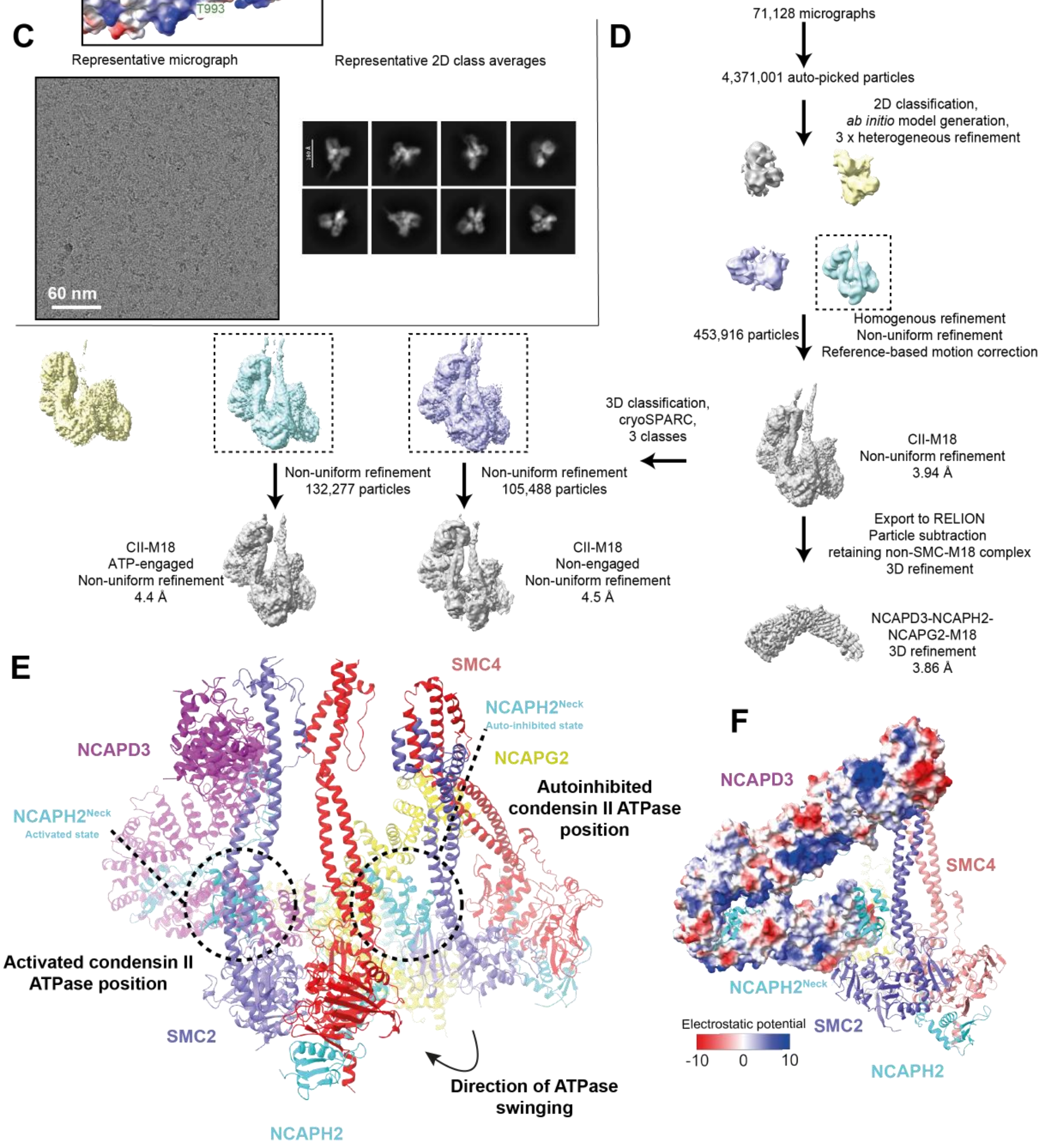

**Figure S7. Mass spectrometry and workflow for cryo-EM reconstructions of phosphorylated condensin II-ATP-M18BP1 (A)** Coulombic potential displayed around the NCAPG2 subunit indicates the presence of positively charged surface residues in proximity to highlighted phosphorylated M18BP1 residues detected by mass spectrometry **(B)** Table listing CDK-phosphorylated residues detected by mass spectrometry within structurally resolved residues of the NCAPD3$^{Tail}$ under a variety of conditions. Relative ratios of phosphorylation between condensin II/M18BP1 and conditions including CDK1 kinase indicate relative enhancement of condensin II phosphorylation by the presence of M18BP1 **(C)** Representative cryo-EM micrograph and 2D class averages arising from vitrified condensin II-M18BP1-ATP complexes **(D)** workflow for the condensin II-M18BP1-ATP/ADP complex data processing **(E).** Structural model showing conformational rearrangement of the SMC2-SMC4 subunits relative to the NCAPD3-NCAPG2 HEAT-repeat subcomplex. Superpositioning was guided by the NCAPD3-NCAPG2 subunits. **(F)** Coulombic potential displayed around the NCAPD3 subunit indicates the presence of a solvent-exposed, positively charged surface.

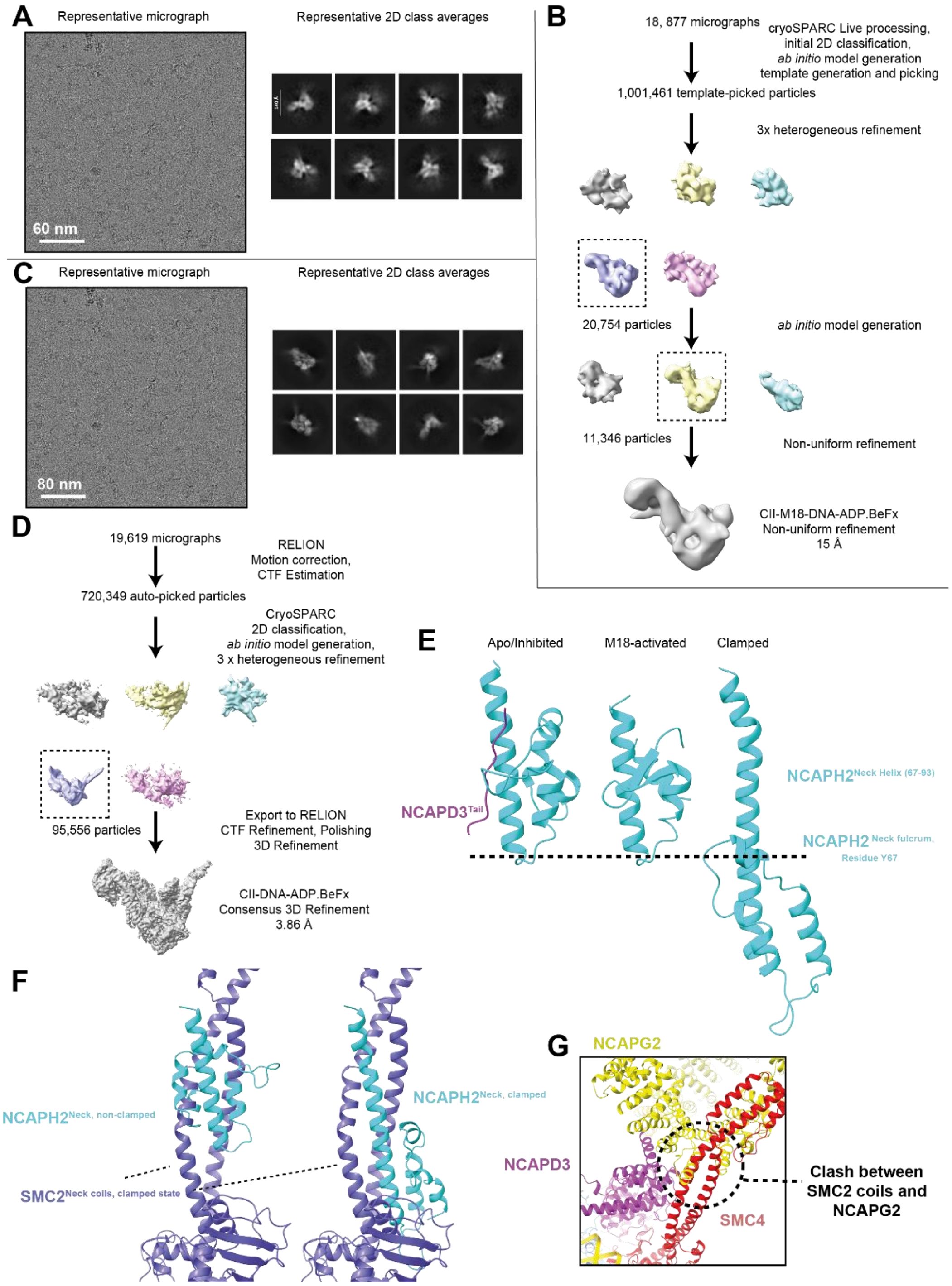
A
Representative micrograph
Representative 2D class averages
60 nm
B
18, 877 micrographs
cryoSPARC Live processing,
initial 2D classification,
ab initio model generation
template generation and picking
1,001,461 template-picked particles
3x heterogeneous refinement
20,754 particles
ab initio model generation
11,346 particles
Non-uniform refinement
CII-M18-DNA-ADP.BeFx
Non-uniform refinement
15 Å
C
Representative micrograph
Representative 2D class averages
80 nm
D
19,619 micrographs
RELION
Motion correction,
CTF Estimation
720,349 auto-picked particles
CryoSPARC
2D classification,
ab initio model generation,
3 x heterogeneous refinement
95,556 particles
Export to RELION
CTF Refinement, Polishing
3D Refinement
CII-DNA-ADP.BeFx
Consensus 3D Refinement
3.86 Å
E
Apo/Inhibited
M18-activated
Clamped
NCAPH2Neck Helix (67-93)
NCAPD3Tail
NCAPH2 Neck fulcrum,
Residue Y67
F
NCAPH2Neck, non-clamped
SMC2Neck coils, clamped state
NCAPH2Neck, clamped
G
NCAPG2
NCAPD3
SMC4
Clash between
SMC2 coils and
NCAPG2

**Figure S8. Workflow for the condensin II- ADP.BeFx-DNA-M18BP1, and condensin II-ADP.BeFx-DNA complex cryo-EM reconstructions, and comparison with other structural states of condensin II (A)** Representative cryo-EM micrograph and 2D class averages of the condensin II- ADP.BeFx-DNA-M18BP1 complex **(B)** Workflow of condensin II- ADP.BeFx-DNA-M18BP1 data processing **(C)** Representative cryo-EM micrograph and 2D class averages of the condensin II- ADP.BeFx-DNA complex **(D)** Workflow of condensin II- ADP.BeFx-DNA data processing **(E)** Structural comparison of the NCAPH2$^{Neck}$ subunit showing extension of the NCAPH2$^{Neck}$ helix upon DNA clamp formation. **(F)** Structural comparison of the NCAPH2$^{Neck}$ between non-clamped and clamped conformations. The N-terminus of the non-clamped NCAPH2$^{Neck}$ folds around the NCAPH2 Y67 fulcrum resulting in structural mimicry of the SMC2$^{Neck\ coils}$ in the clamped state. Superpositioning was guided by residues 67-93 of the NCAPH2$^{Neck}$ helix. **(G)** Superpositioning of the NCAPG2-NCAPD3 heterodimer onto the position of NCAPD3 in the clamped state results in a steric clash between the SMC4$^{Coils}$ and NCAPG2.

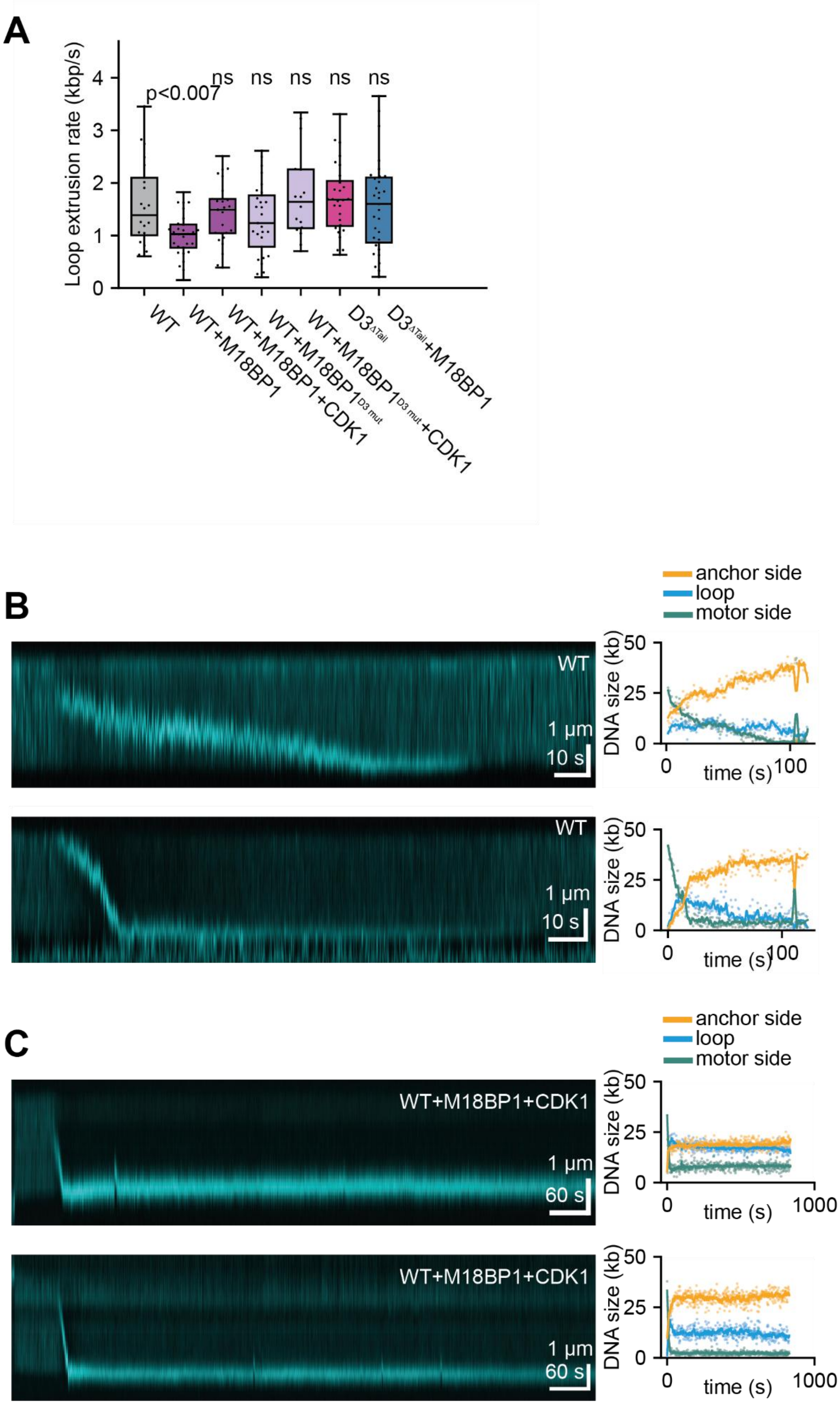
A
Loop extrusion rate (kbp/s)
0
1
2
3
4
p<0.007
ns
ns
ns
ns
ns
WT
WT+M18BP1
WT+M18BP1+CDK1
WT+M18BP1D3 mut
WT+M18BP1D3 mut+CDK1
D3ΔTail
D3ΔTail+M18BP1
B
anchor side
loop
motor side
WT
1 µm
10 s
DNA size (kb)
50
25
0
0
time (s)
100
WT
1 µm
10 s
C
anchor side
loop
motor side
WT+M18BP1+CDK1
1 µm
60 s
DNA size (kb)
time (s)
1000
WT+M18BP1+CDK1
1 µm
60 s

**Figure S9. M18BP1 increases condensin II-loop duration by stabilizing slipping from the anchor side (A)** Box-whisker-plots of loop extrusion rate at loop initiation of loops formed by condensin II WT and in the presence of M18BP1; M18BP1 plus CDK1, M18BP1$^{D3\ mut}$ and M18BP1$^{D3\ mut}$ plus CDK1 and by condensin II D3$^{\Delta Tail}$ and in the presence of M18BP1. The central line denotes the median, the box limit denotes the 25th–75th percentile and the whiskers denote minimum and maxima. The P values were calculated using Welch's t test. **(B, C)** Further examples of representative kymographs of loop extrusion events by WT condensin II (B) and by condensin II in the presence of M18BP1 and Cyclin B/CDK1 (C), showing SxO-stained DNA molecules (left). DNA lengths extracted from the corresponding kymographs in (B) and (C) for regions outside the loop (motor and anchor sides) and within the loop (Loop) (right).

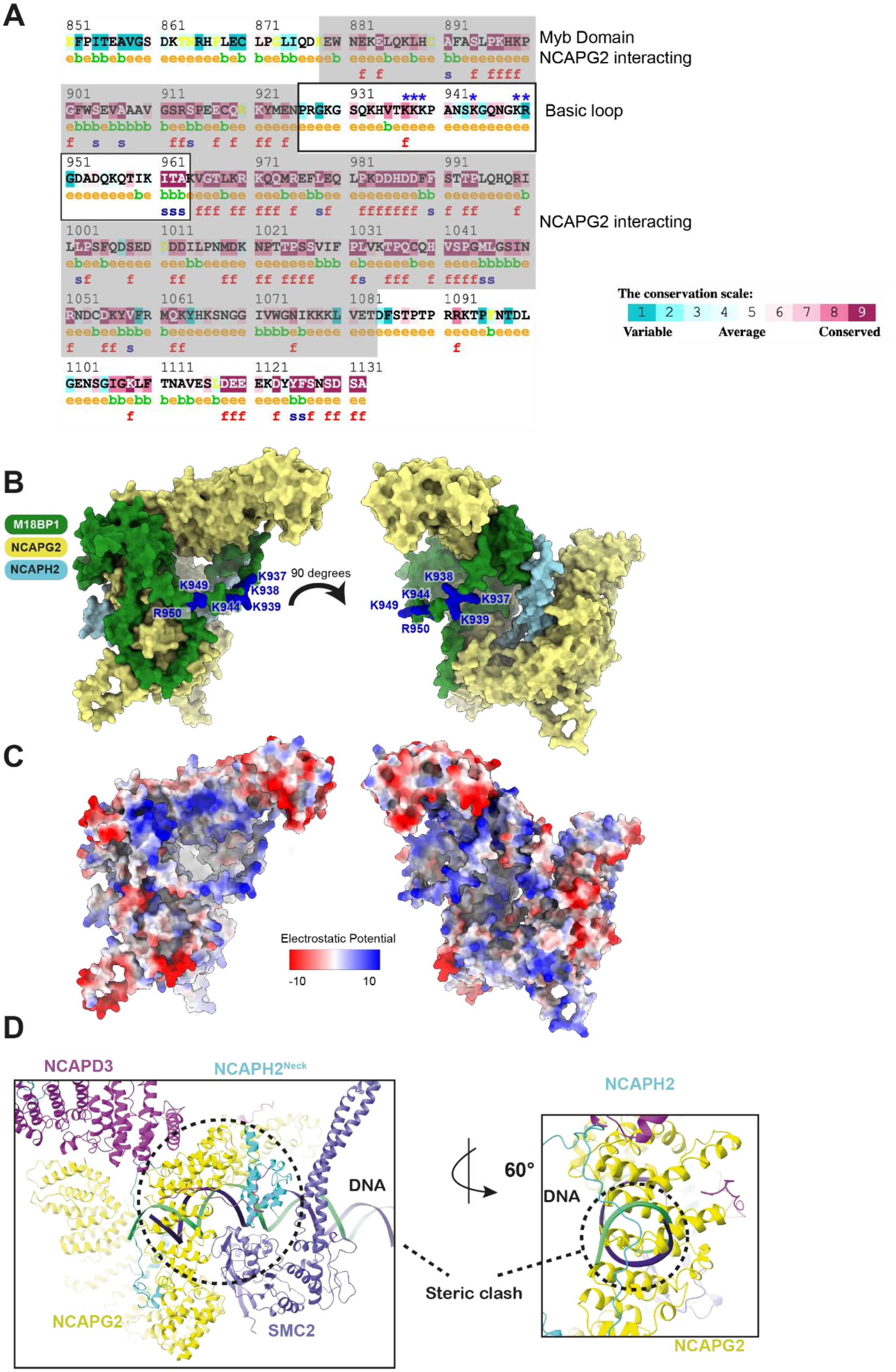
A
Myb Domain
NCAPG2 interacting
Basic loop
NCAPG2 interacting
The conservation scale:
Variable
Average
Conserved
B
M18BP1
NCAPG2
NCAPH2
90 degrees
K937
K938
K939
K944
K949
R950
C
Electrostatic Potential
-10
10
D
NCAPD3
NCAPH2Neck
DNA
NCAPG2
SMC2
60°
NCAPH2
Steric clash

**Figure S10. M18BP1 has a positively charged loop. Further structural comparison of ATP-engaged and ADP.BeFx-DNA-bound condensin II (A)** Consurf[68] sequence conservation indicating conserved positively charged residues. **(B)** Position and **(C)** coulombic potential of Alpha-Fold2 [83] predicted model of NCAPG2-NCAPH2-M18BP1 complex. Basic residues mutated to glutamic acid are indicated with * in (A) and coloured blue in (B). **(D)** Structural comparison of the position of DNA on SMC2 in the clamped state produces a steric clash with the NCAPG2-SMC2 interface in the ATP-engaged state.

Table 1. **Cryo-EM data collection, refinement and validation statistics**

| | Apo condensin II protomer monomer<br><br>PDB 28PG / EMD-56706 | ATP-engaged condensin II<br><br>PDB 28OZ / EMD-56696 | ATP-engaged condensin II – M18BP1<br>PDB 28ML / EMD-56616 | Condensin II – DNA – ADP.BeFx<br><br>PDB: 28IA / EMD-56535 |
|---|---|---|---|---|
| **Data collection and processing** | | | | |
| Magnification | 130,000 | 165,000 | 165,000 | 130,000 |
| Voltage (kV) | 300 | 200 | 200 | 300 |
| Electron exposure (e–/Å$^2$) | 40 | 50 | 50 | 40 |
| Defocus range (μm) | 0.8-2.2 | 0.8-2.2 | 0.8-2.2 | 0.8-2.2 |
| Pixel size (Å) | 0.921 | 0.69 | 0.69 | 0.921 |
| Symmetry imposed | C1 | C1 | C1 | C1 |
| Initial particle images (no.) | 1,822,120 | 2,249,000 | 4,371,001 | 720,349 |
| Final particle images (no.) | 300,224 | 270,728 | 132,277 | 95,556 |
| Map resolution (Å) | 3.9 | 3.8 | 4.39 | 3.86 |
| FSC threshold | 0.143 | 0.143 | 0.143 | 0.143 |
| Map resolution range (Å) | 3.9-20 | 3.8-20 | 4.39-20 | 3.86-20 |
| **Refinement** | | | | |
| Initial model used (PDB code) | *Ab initio*, AlphaFold3 | *Ab initio*, AlphaFold3 | *Ab initio*, AlphaFold3 | *Ab initio*, AlphaFold3 |
| Model resolution (Å) | 3.9 | 3.8 | 4.39 | 3.86 |
| FSC threshold | 0.143 | 0.143 | 0.143 | 0.143 |
| Model resolution range (Å) | 3.9-20 | 3.8-20 | 4.39-20 | 3.86-20 |
| Map sharpening *B* factor (Å$^2$) | -88 | -88 | -88 | -88 |
| Model composition | | | | |
| Non-hydrogen | 17406 | 39620 | 53989 | 21366 |
| atoms | 2184 | 3738 | 3845 | 2473 |
| Protein residues | 0 | 0 | 0 | 72 |
| Nucleotide | | | | |
| Ligand | 0 | 4 | 0 | 6 |
| *B* factors (Å$^2$) | | | | |
| Protein | 125.85 | 373.57 | 325.45 | 128.82 |
| Nucleotide | N/A | N/A | N/A | 229.27 |
| Ligand | N/A | 153.73 | N/A | 33.82 |
| R.m.s. deviations | | | | |
| Bond lengths (Å) | 0.002 | 0.010 | 0.002 | 0.005 |
| Bond angles (°) | 0.494 | 0.601 | 0.545 | 0.836 |

| | | | | |
|---|---|---|---|---|
| Validation | | | | |
| MolProbity score | 1.30 | 2.27 | 2.06 | 2.1 |
| Clashscore | 5.47 | 14.21 | 14.35 | 21.74 |
| Poor rotamers (%) | 0.00 | 0.24 | 1.96 | 0.82 |
| Ramachandran plot | | | | 5 |
| Favored (%) | 97.97 | 96.78 | 96.98 | 96.12 |
| Allowed (%) | 2.03 | 2.98 | 2.97 | 3.88 |
| Disallowed (%) | 0.00 | 0.24 | 0.05 | 0.00 |

#### Table 2. DNA constructs.

| Recombinant DNA | Reference |
|---|---|
| pF1 NCAPG2-NCAPD3 | This study |
| pF1 NCAPH2-TEV-2xStrepII | This study |
| pF1 SMC2-SMC4 | This study |
| pF1 NCAPG2-NCAPD3 | This study |
| pF1 NCAPG2-NCAPH2 313-415 | This study |
| pBIG2abc Condensin II strep | Kong et al [19] |
| pBIG2abc Condensin II ybbR strep | Kong et al [19]. |
| pBIG2abc Condensin IIΔD3 strep (Lacking NCAPD3) | Houlard et al[17] |
| pBIG2abc Condensin IIΔG2 strep (Lacking NCAPG2) | Houlard et al[17] |
| pBIG2abc Condensin II QL ΔD3 strep (Lacking NCAPD3, SMC2 Q147L, SMC4 Q229L) | This study |
| pBIG2abc Condensin II QL ΔG2 strep (Lacking NCAPG2, SMC2 Q147L, SMC4 Q229L) | This study |
| pLIB NCAPD3 ΔTail (Deletion of NCAPD3 1298-1498) | This study |
| pLIB NCAPD3 ΔTail-G2 (Deletion of NCAPD3 1470-1475) | This study |
| NCAPD3 Tail-H2ala (NCAPD3 1453-1458, inclusive, to Alanine) | |
| pET 10xHis M18BP1 874-1132 ybbR MBP | This study |
| pET 10xHis M18BP1 874-1132 ybbR MBP D3 mut (W1073A, I1076A, K1077E, K1078E) | This study |
| pET M18BP1 NC (K937E, K938E, K939E, K944E, K949E, R950E) | This study |
